# The endophytic fungus *Cosmosporella* sp. VM-42 from *Vinca minor* is a source of bioactive compounds with potent activity against drug-resistant bacteria

**DOI:** 10.1101/2024.12.27.630490

**Authors:** Ting He, Xiao Li, Rosario del Carmen Flores-Vallejo, Ana-Maria Radu, Jan Maarten van Dijl, Kristina Haslinger

## Abstract

Medicinal plants serve as valuable resources for the isolation of endophytic fungi. *Vinca minor* is a well-known producer of important vinca alkaloids and emerges as a promising source of endophytic fungi with antibacterial potential and biosynthetic capacity. In this study, we isolated an endophytic fungus from *V. minor* and identified it as *Cosmosporella* sp. VM-42. To date, relatively little is known about this fungal genus. The ethyl acetate extract of this isolate selectively inhibited Gram-positive bacteria, such as methicillin-sensitive and methicillin-resistant *Staphylococcus aureus* (MSSA and MRSA). Therefore, we isolated the most abundant compound from the crude extract and identified it as nectriapyrone with MIC and MBC values ranging from 125 to 62.5 µg/mL against MSSA and MRSA strains. We further sequenced and annotated the 39.07 Mb genome of the isolate, revealing that it encodes 9,842 protein-coding genes, including 415 genes for carbohydrate-active enzymes and various biosynthetic gene clusters. Our untargeted metabolomic analysis shows that the fungus produces various secondary metabolites, including cyclodepsipeptides, dimeric naphtho-γ-pyrones, and macrolactones, which are known to have antifungal and antibacterial activities. In addition, we used small-molecule epigenetic modulators to activate the expression of silent biosynthetic gene clusters to broaden the chemical profile of *Cosmosporella* sp. VM-42. Taken together, we provide a first systematic analysis of *Cosmosporella* sp. VM-42, and our results show that it is a promising source of compounds with pharmacological potential against drug resistant bacteria.

**Highlights:**

- First comprehensive study of *Cosmosporella* sp. VM-42 by genomics and metabolomics
- The fungus produces chemically-diverse secondary metabolites with antibacterial activity
- Nectriapyrone, the main compound, shows bactericidal activity against MSSA and MRSA
- Small-molecule epigenetic modulators trigger production of putatively new secondary metabolites.
- Other secondary metabolites of *Cosmosporella* may present novel bioactivities.

## 1. Introduction

The emergence of drug-resistant pathogenic microorganisms has developed into a serious threat to human health worldwide (WHO, 2022). Many of the existing drugs used to treat infectious diseases have become ineffective (Fisher et al., 2022; Lee Ventola, 2015). Thus, the quest for finding novel drugs of natural origin to alleviate this pressure has become the utmost priority. Many plants and microorganisms are important sources of bioactive compounds. In past decades, also endophytic fungi associated with medicinal plants have received attention as possible producers of novel bioactive compounds (Qi et al., 2019; Wu et al., 2018). Such fungi exist in a symbiotic relationship with their host, and have been shown to produce secondary metabolites that contribute to the fitness and survival of the plant (Baron and Rigobelo, 2022). These bioactive compounds present unique chemistry and interesting biological activities against bacteria (Augner et al., 2013; Kjer et al., 2009), fungi (Pereira et al., 2015), or cancers (Monroe E. Wall et al., 1966; Stierle et al., 1993).

Approximately 300,000 known species of higher plants exist on Earth, each serving as a host to one or more species of endophytic fungi (Strobel and Daisy, 2003). It is estimated that the number of endophytic fungi alone could approach one million. Yet only 5% of fungal species have been described to date (Bhunjun et al., 2023). Despite initiatives to sequence more fungal genomes, such as the 1000 Fungal Genomes Project at the Joint Genome Institute, endophytic fungi remain a relatively under-investigated group of microorganisms, with much of their potential yet to be explored (Sagita et al., 2021). The number of pharmacologically important compounds (Deshmukh et al., 2015; Wen et al., 2022) and valuable enzymes (Corrêa et al., 2014; Srinivas et al., 2013) identified in endophytic fungi continuously increases, driven by advances in the combination of several “omics” approaches. However, the exploration of endophytic fungi as new sources of bioactive compounds and enzymes is challenged by the silencing of biosynthetic gene clusters under laboratory conditions. Various strategies, including One Strain Many Compounds, co-culture, and chemical epigenetic modification have been investigated to address these issues (Hewage et al., 2014; H. T. Li et al., 2019).

*Vinca minor*, is a medicinal plant of the *Apocynaceae* family, which is known to produce important vinca alkaloids. In folk medicine, it is used internally for circulatory disorders, cerebral circulatory impairment and brain metabolism support (LaGow B, 2004). Interestingly, the endophytic microbiota of *V. minor* has remained largely unexplored. In the present study, we applied: 1) *in vitro* antibacterial bioassays against drug-sensitive and -resistant bacteria; 2) untargeted metabolomics analysis comparing plain fungal cultures and cultures treated with small molecule epigenetic modulators; and 3) whole genome sequencing and genome mining, to unravel the antibacterial and biosynthetic potential of an endophytic fungus belonging to the *Cosmosporella* genus, which was isolated from *V. minor*. Overall, we present the first comprehensive exploration of a *Cosmosporella* isolate, showing that this fungus is a source of bioactive compounds with potential pharmacological applications against drug-resistant bacteria.

## 2. Materials and Methods

### 2.1. Fungus isolation and cultivation

The *Cosmosporella* sp. VM-42 strain was isolated from healthy-looking, surface-sterilized leaves of *V. minor* as described previously (He et al., 2023). Briefly, leaves of *V. minor* were freshly collected in Groningen (The Netherlands) in November 2021, washed in an ultrasonic water bath (160 W, 15 min), surface-sterilized in 70% ethanol for 1 min, followed by 1% sodium hypochlorite for 2 min, then washed in distilled water for 3 × 1 min. Leaves were aseptically cut into small fragments and directly placed on potato dextrose agar (PDA) medium, supplemented with 100 mg·L^−1^ ampicillin and 30 mg·L^−1^ kanamycin, and incubated at 25°C for 2 to 4 weeks. 200 µL water from the last washing step was also inoculated onto PDA and incubated at 25°C to check the effectiveness of surface sterilization. Fungal mycelium emerging from leaf pieces was collected and purified by re-streaking on fresh PDA. Plates with purified colonies were sealed with parafilm and stored at 4°C.

### 2.2. Morphological analysis, Internal Transcribed Spacer (ITS)-based identification, and multilocus phylogeny of the Cosmosporella sp. VM-42

For morphological characterization, the fungal isolate was grown on Malt Extract Agar (MEA), Czapek’s Agar (CZA), Sabouraud dextrose agar (SDA), Czapek yeast autolysate agar (CYA), and PDA. After 28 days at 25°C, the front and back colour, and diameter of the fungal colonies were recorded. For microscopic analysis, mycelium samples were collected and mixed with lactophenol blue dye or water and visualized using an optical microscope (Olympus BX41) with a Leica camera (Heerbrugg, Switzerland).

Internal Transcribed Spacer (ITS) sequencing performed as previously described (He et al., 2023). Briefly, the full ITS region was amplified by Polymerase Chain Reaction (PCR) with the ITSF1 and ITS4 primers (Gardes and Bruns, 1993) in a thermal cycler with a Q5 PCR master mix (New England Biolabs, Ipswich, MA, United States) and fungal DNA using the following program: 1 min 98°C, 30 cycles of 10 s 98°C, 15 s 55°C, 20 s 72°C, followed by 5 min of final extension at 72°C. After Sanger sequencing (Macrogen, Amsterdam, Netherlands), the sequences were analysed with the Basic Local Alignment Search Tool (BLAST) against the nucleotide collection of the National Center for Biotechnology Information (NCBI) to identify the best match for the fungal isolate based on E-value.

For multilocus phylogenetic analysis, three barcode sequences were extracted from the whole genome assembly (vide infra): the genes coding for beta-tubulin (*TUB2*), as well as the sequence of the ITS and the large ribosomal subunit (LSU). The phylogenetic tree was built on sequences of 23 taxa in the *Nectriaceae* family, downloaded from NCBI (*Fusarium oxysporum* Fo47 as an outgroup, Table S1). Sequences from the three loci were concatenated (∼ 1500 nucleotides in total) and aligned using ClustalW, followed by the generation of a Maximum-Likelihood (ML) tree in MEGA (version 11) (Tamura et al., 2021) with 1000 bootstrap replicates.

2.3. *Fermentation, extraction, isolation, and structural identification of isolated compounds*

Five mycelial discs of *Cosmosporella* sp. VM-42 grown on PDA plates at 25℃ for 14 days were used to inoculate two Erlenmeyer flasks, each of them containing 1 L sterile PDB medium (potato dextrose broth powder 24 g/ L). The flasks were incubated at 25℃ for 14 days at 100 rpm. The fermentation broth and mycelium were separated by filtration and extracted with 1 L and 3 L of organic solvent (EtOAc-MeOH-formic acid in the ratio of 900: 100: 1) using liquid-liquid extraction and ultrasonic extraction, respectively. The organic phases of both extractions were combined and concentrated under reduced pressure (Hei-VAP Core Rotary Evaporators, Heidolph Instruments GmbH & CO. KG, Germany) at 4℃ and freeze-dried (Lyovapor☐ L-200 Basic, BÜCHI Labortechnik, Switzerland) to yield a total of 4.34 g crude extract.

For antibacterial assays a soluble ethyl acetate fraction (sEtOAc) was obtained using 200 mg of crude extract resuspended with 20 mL EtOAc. This mixture was sonicated for 20 min using a Branson 5800® water bath sonicator. Following centrifugation for 20 min (8000 g’s and 4°C), the sEtOAc fraction was recovered in a glass scintillation vial. This process was repeated twice with the remaining insoluble fraction mixed with other 20 mL EtOAc. The sEtOAc fractions were pooled together and dried under gentle stream of N_2_.

For compound isolation, a total of 1069 mg crude extract was dissolved in 15 mL MilliQ-water and centrifuged to obtain the supernatant for fractionation by flash chromatography. The supernatant was loaded onto a C18 column (FlashPure EcoFlex C18 12 g column from Buchi®) in a BÜCHI Reveleris™ X2-UV System, and eluted with a 20% ACN (solvent A)-ACN (solvent B) gradient program (10 min, 0% B; 10 min, 20% B; 10 min, 40% B; 10 min, 60% B; 10 min, 80% B; 5 min, 95% B; 10 mL/min, UV detection at 220, 270, and 330 nm) to give 37 fractions. Fractions 19 to 27, which contained the target compound based on LR-LC-MS analysis, were pooled together and concentrated under reduced pressure at 40℃. The concentrate was loaded onto the same C18 column and eluted with H_2_O (A)-ACN (B) in a gradient program (5 min, 10% B; 5 min, 30% B; 5 min, 40% B; 5 min, 45% B; 6 min, 46% B; 4 min, 75% B; 4 min, 95% B). After checking the fractions by LR-HPLC-MS, those fractions mainly consisting of the target compound with m/z 195 were pooled and freeze-dried to yield 3 mg of compound **27**. The isolated compound was stored at –20℃ until further use in antibacterial assays and structural analysis by nuclear magnetic resonance spectroscopy (NMR).

For NMR analysis 3 mg of compound **27** was dissolved in 550 µL of DMSO-d6 .^1^H (recorded at 600 MHz in DMSO-d6), ^13^C (recorded at 150 MHz in DMSO-d6), and two-dimensional (2D) NMR (^1^H-^1^H COSY, HSQC, and HMBC) analysis was carried out on a Bruker Ascend Evo TM 600 NMR spectrometer (Bruker-Biospin, Billerica, MA, USA). MestReNova 14 was used for data analysis.

### 2.4. Antibacterial susceptibility testing

We initially screened for antibacterial activity of the *Cosmosporella* sp. VM-42 using the agar diffusion method against the non-pathogenic indicator bacterium *Bacillus subtilis* (Balouiri et al., 2016). Briefly, we inoculated the fungus on PDA and incubated it for 5 days at 25°C. Afterwards, 250 µL of bacterial suspension with optical density at λ = 600 nm (OD_600_) of 0.1 was inoculated on top of the fungal colony and incubated overnight at 37°C. Inhibition of the bacterial mat growth around the fungal colony was noted by visual inspection after the incubation time.

Later, we analyzed the sEtOAc fraction and compound **27** with the broth microdilution method considering the guidelines of the European Committee on Antimicrobial Susceptibility Testing. The test substances were completely dissolved in dimethyl sulfoxide (DMSO), filter-sterilized with 0.22 µm PTFE membranes and stored at 4°C, protected from light until used. Six in-test concentrations were evaluated (15.625 - 500 µg/ mL) against *Staphylococcus aureus* strains (methicillin-sensitive (MSSA) S.a. ATCC 29213, S.a. NCTC 8325, S.a. Newman, and S.a. HG001 and methicillin-resistant (MRSA) S.a. USA 300) and *Escherichia coli* ATCC 25922. Bacteria were grown on Blood Agar 2 (BA2) for 24 h at 36 ± 1°C. Then an overnight culture was prepared in Mueller Hinton (MH) broth incubated at 36 ± 1°C with constant shaking at 250 rpm. Each bacterial culture was seeded into a 96-well microtiter plate at a final concentration of 5 × 10^4^ colony-forming units (CFUs) per well. The plate was incubated at 36 ± 1°C for 24 h with continuous double orbital shaking (282 rpm) inside a Biotek Epoch-2 ® microplate reader. The MIC was defined as the lowest concentration of the extract or compound that completely inhibited bacterial growth (complete absence of haze growth, pinpoint growth, or turbidity). The final concentration of DMSO in the assay was ≤ 1% (v/ v) and served as a negative control. In addition, bacteria treated with a sterile solution of 10% (v/ v) of DMSO in Milli-Q water and sterile Milli-Q water served as positive and negative controls, respectively. To quantify the results of the antibacterial assay, OD_600_ values were recorded at the beginning of the incubation (t = 0 h) and at the end of the incubation time (t = 24 h) with the microplate reader. The percentage inhibition of bacterial growth was calculated as: Bacterial growth (%) = (mean OD_600_ value *fraction or compound-treated bacteria at t_24h_* - mean OD_600_ value *fraction or compound-treated bacteria at t_0h_*) / (mean OD_600_ value of *vehicle-treated bacteria t_24h_* - mean OD_600_ value of *vehicle-treated bacteria t_0h_*) ×100. Bacteria treated only with Milli-Q water were considered as reference for establishing 100% of bacterial growth.

Additionally, for compound **27,** the Minimum Bactericidal Concentration (MBC) was determined using the same microplate that was used for the MIC determination. Bacterial broth from the microplate wells treated with the compound at concentrations at or above the MIC was homogenized by pipetting before spotting 2.5 µL on MH agar plates for incubation at 36 ± 1°C for 24 h. The MBC was determined by visual inspection and counting bacterial CFUs. The MBC was defined as the lowest concentration of the compound at which approximately 99.99% of the bacterial population was killed causing inhibition of colony formation after incubation. Data analysis was performed in GraphPad Prism v.8.0.1.

### 2.5. Untargeted metabolomics

#### 2.5.1. Sample preparation and data acquisition

Fungal mycelium was transferred to small (⌀ 35 mm × 10 mm) PDA plates supplemented with different concentrations (0, 1, 10, and 100 mM) of the histone deacetylase (HDAC) inhibitors sodium butyrate (SB) or procaine hydrochloride (PH). The plates were incubated at 25°C for 14 days alongside uninoculated PDA, PDA-SB, and PDA-PH plates as controls.

For extraction of secondary metabolites (SMs), the whole agar pads including mycelium were sliced and transferred to 25 mL glass bottles, then mixed with 4 mL solvent (9:1 ethyl acetate-methanol (*v*/*v*) with 0.1% formic acid), spiked with 5 μL caffeine standard solution (10 mg/ mL stock), and sonicated in a sonication bath for one hour. The organic phase was collected and dried under a gentle stream of N_2_. Dried extracts were resuspended in 500 μL 1:1 MeOH-Milli-Q water (*v*/*v*) and filtered with 0.45 μm PTFE filters.

Low-Resolution Liquid Chromatography-coupled Mass Spectrometry (LR-LC-MS) was performed with a Waters high-performance liquid chromatography (HPLC) system controlled by MassLynx v 4.2 and equipped with a Waters ACQUITY Qda Mass Detector, and a Waters 2998 PDA detector, using a Waters XBridge C18 analytical column (2.1 × 50 mm, 3.5 μm). MilliQ water with 0.1% formic acid (FA) (phase A) and ACN with 0.1% FA (phase B) were used as solvents at a flow rate of 0.25 mL/min with the following program: 0-2 min, 95% A; 2-17 min, linear decrease to 50% A; 17-21 min, linear decrease to 10% A; 21-24 min, 10% A; 24-24.01 min, increase to 95% A; 24.01-30 min, 95% A. The injection volume was 2 μL, and the column temperature 40°C. MS analysis was performed in positive and negative electrospray ionization (ESI) mode. The mass data was acquired with a survey scan at *m*/*z* 100-1250.

High-Resolution Liquid Chromatography-coupled tandem Mass Spectrometry (HR-LC-MS/MS) was performed with a Shimadzu Nexera X2 HPLC system with binary LC20ADXR pump coupled to a Q Exactive Plus hybrid quadrupole-orbitrap mass spectrometer (Thermo Fisher Scientific, Waltham, MA, USA). A Kinetex EVO C18 reversed-phase column (100 mm × 2.1 mm I.D., 2.6 μm, 100 Å particles, Phenomenex, Torrance, CA, United States) maintained at 50°C was used for separations. The mobile phase consisted of solution A (0.1% formic acid in Milli-Q water) and solution B (0.1% formic acid in Acetonitrile). A linear gradient was used: 0–2 min 5% B, 2–17 min linear increase to 50% B, 17–21 min linear increase to 90% B, 21–24 min 90% B, 24–24.01 min decrease to 5% B, and 24.01–30 min 5% B. The injection volume was 2 µL, and the flow rate was set to 0.25 mL/min. MS and MS/MS analyses were performed with ESI in positive mode at a spray voltage of 3.5 kV and sheath and auxiliary gas flow set at 60 and 11, respectively. The ion transfer tube temperature was 300°C. Spectra were acquired in data-dependent mode with a survey scan at *m/z* 100-1500 at a resolution of 70,000, followed by MS/MS fragmentation of the top 5 precursor ions at a resolution of 17,500. A normalized collision energy of 30 was used for fragmentation, and fragmented precursor ions were dynamically excluded for 10 s.

#### 2.5.2. Data analysis

The acquired data were further processed by Thermo Scientific FreeStyle software version 1.8. The raw MS/MS data file was converted to mzXML format using the easy convertor provided by the Global Natural Products Social Molecular Networking (GNPS) (https://ccms-ucsd.github.io/GNPSDocumentation/fileconversion/). The data files were subsequently uploaded to GNPS (https://gnps.ucsd.edu/) using WinSCP.

A molecular network was created using the online workflow on the GNPS website. The data were filtered by removing all MS/MS fragment ions within ±17 Da of the precursor *m/z*. MS/MS spectra were window filtered by choosing only the top six fragment ions in the ±50 Da window throughout the spectrum. The precursor ion mass tolerance and MS/MS fragment ion tolerance were both set to 0.02 Da. A network was then created where edges were filtered to have a cosine score above 0.7 and more than six matched peaks. Further, edges between two nodes were kept in the network if each of the nodes appeared in the other’s respective top 10 most similar nodes (molecular networking job: https://gnps.ucsd.edu/ProteoSAFe/status.jsp?task=6811c4d86d9f4c289e60a5a303ad930d). All mass spectrometry data were deposited on GNPS under the accession number MassIVE ID: MSV000095645. The molecular network was visualized in Cytoscape version 3.9.1 (Otasek et al., 2019). Nodes that also existed in the PDA-SB and PDA-PH controls were considered background and thus omitted from the final molecular network.

The network file .graphML was downloaded from GNPS and further uploaded to SNAP-MS (https://www.npatlas.org/discover/snapms/)(Morehouse et al., 2023). NPAtlas Fungi was selected as reference database and other options were default. The result was visualized in Cytoscape version 3.9.1Click or tap here to enter text. and data can be found in supplementary material.The raw MS/MS data files were also imported into Mzmine 4.0.3 (Schmid et al., 2023). After setting the processing wizard with the default parameters, an .mgf file for SIRIUS analysis was obtained and imported to SIRIUS 5.8.6 (Dührkop et al., 2019) (https://boecker-lab.github.io/docs.sirius.github.io/). Files can be found in supplementary material. Peaks were subsequently classified and annotated by SIRIUS-molecular formula identification, ZODIAC (Ludwig et al., 2020), CSI:FingerID (Dührkop et al., 2015), and CANOPUS (Djoumbou Feunang et al., 2016; Dührkop et al., 2021) under the default parameter settings, in addition to the instrument parameter “Orbitrap” in molecular formula identification option and the selected fallback adducts in CSI:FingerID, including [M + H]^+^, [M - H_2_O + H]^+^, [M + H_3_N + H]^+^, [M + H_2_O + H]^+^, [M + K]^+^, and [M + Na]^+^. The molecular formulae were determined when SIRIUS scores exceeded 95%, and the compound structures were determined when similarity was higher than 70%. The output of GNPS, SNAP-MS, and SIRIUS was combined to obtain a reliable compound profile.

### 2.6. Genome mining

#### 2.6.1. Whole genome sequencing and assembly

DNA extraction was performed according to previously described methods (He et al., 2023). Briefly, the fungal isolate was grown in 25 mL Dextrose Peptone Yeast medium at 25°C for 5 days before the mycelium was collected, flash-frozen in liquid nitrogen, lyophilized overnight using Lyovapor™ L-200, and stored at −20°C.

DNA was extracted using QIAGEN Genomic-Tips 20 G-1 and the QIAGEN Genomic Buffer Set according to the manufacturer’s protocol with the following minor modifications: (1) Six 2 mL Eppendorf tubes with 25 mg lyophilized and ground mycelium were used instead of cells collected directly from the medium; (2) Vinotaste PRO (Novozymes, Bagsværd, Denmark) with a final concentration of 20 mg/mL was used as lytic enzyme instead of lyticase; (3) Enzymatic degradation of the cell wall was performed at 30°C for 1 h (100 rpm) and cell lysis was performed at 50°C for 2 h (25 rpm) instead of the recommended time and temperature; (4) QIAGEN Genomic-Tips 20 G-1 were washed four times.

Circulomics Short Read Eliminator XS (PacBio, Menlo Park, CA, United States) was used to remove small fragments. Quality control of the purified DNA was performed using a NanoDrop N-100 (ThermoFisher, Waltham, MA, United States) and the Qubit 3.0 dsDNA HS Assay Kit (Invitrogen, Waltham, MA, United States).

For long-read sequencing, the genomic DNA (Table S2) was prepared using Oxford Nanopore Technologies’ Ligation sequencing kit V12 (SQK-LSK112) according to the manufacturer’s guidelines. Briefly, genomic DNA (1000 ng) was subjected to end repair and tailing by NEBNext FFPE DNA Repair mix and NEBNext Ultra II End repair/dA-tailing modules (New England Biolabs, Ipswich, MA, United States) and purified with an equal volume of AMPure XP (Beckman Coulter, Pasadena, CA, United States) magnetic beads. The sequencing adaptors were ligated using the NEBNext Quick Ligation Module (New England Biolabs, Ipswich, MA, United States). The mix was further purified with 0.4 × volume of AMPure XP Beads and cleaned up using the Long Fragment Buffer. The sequencing library was loaded into a primed FLO-MIN112 (ID: FAT75549) flow cell on a MinION device. Data acquisition and real-time basecalling were carried out with MinKNOW software (version 22.05.5).

The raw reads were basecalled using Guppy version 6.1.5 (Oxford Nanopore Technologies, Oxford, UK) in GPU mode using the dna_r10.4_e8.1_sup.cfg model (Wick et al., 2019). The basecalled reads were filtered to a minimum length of 2 kb and a minimum quality of Q10 using NanoFilt (version 2.8.0) (Coster et al., 2018). NanoPlot (version 1.40.0) (Coster et al., 2018) was used to evaluate the filtered reads. Assembly was performed using Flye (version 2.9-b1778). The quality of the genome assembly was evaluated using QUAST v5.1.0rc1 (Gurevich et al., 2013). Bandage (version 0.8.1) (Wick et al., 2015) was used to visualize the newly assembled genome of *Cosmosporella* sp. VM-42 (Figure S1). The draft assembly was subsequently polished in two rounds: first using Racon version 1.4.10 with default settings (Vaser et al., 2017), then Medaka version 0.11.5 with default settings. The completeness of the assembly was evaluated using BUSCO 5.4.3 (ascomycota_odb10 dataset). Genome annotation was carried out using the online platform Genome Sequence Annotation Server (GenSAS, https://www.gensas.org, accessed on 5 September 2023), which provides a pipeline for whole-genome structural and functional annotation (Humann et al., 2019). We also assembled the mitochondrial genome using 10,000 reads and annotated it with the web-based tool GeSeq (Tillich et al., 2017). The sequencing data and genome assembly for this study were deposited in the European Nucleotide Archive (ENA) at EMBL-EBI under accessions ERR14081000 and ERZ24985901, respectively.

#### 2.6.2 Gene prediction and annotation

The tRNA and rRNA genes were predicted using tRNA scan-SE (version 2.0.11) (Chan et al., 2021) and barrnap (version 0.9) (Gurevich et al., 2013). Gene Ontology (GO) annotation was performed using Pannzer2 (Törönen and Holm, 2022). To predict CAZymes, we used the web-based meta server dbCAN2 (http://cys.bios. niu.edu/dbCAN2), which integrates three tools (dbCAN HMM, CAZy, and dbCAN-sub) (Zhang et al., 2018). The three outputs were combined, and CAZymes found by only one tool were removed to improve the CAZyme annotation accuracy. Secondary metabolite biosynthetic clusters were identified using the antiSMASH web server (fungal version 7.0) with the default settings (Harwood et al., 2023).

## 3. Results

### 3.1. Isolate VM-42 from Vinca minor is a Cosmosporella species

To learn more about the endophytic fungi residing within *Vinca minor*, we isolated 50 fungi from healthy leaves of *V. minor* collected in Groningen, The Netherlands. We then screened the antibacterial activities of these endophytes against Gram-positive and Gram-negative bacteria with an agar diffusion method. One of the endophytic fungi exhibited a strong inhibitory effect against *Bacillus subtilis* showing an inhibition zone with a diameter of 35-40 mm (Figure S2A). We preliminarily classified this isolate based on ITS sequencing and comparison with the NCBI database as a close relative of *Cosmosporella* and *Cosmospora* species. Since neither whole-genome sequences nor metabolomic analyses were publicly available for these genera, we decided to systematically explore this fungus. We performed long-read sequencing of genomic DNA with the Oxford Nanopore Technology (ONT), yielding 1.09 million raw reads (6.8 Gbp) with an N50 of 10.04 kb and high read quality of 19.2 after filtering (Table 1). Subsequently, we assembled the filtered reads into 10 scaffolds with a total size of 39.07 Mbp (∼120 × coverage, Figure S1). The polished draft assembly has a BUSCO completeness of 97.3%, indicating a highly contiguous and complete genome sequence. Next, we structurally annotated the assembly with GenSAS and predicted 11,642 protein-encoding genes, 55 rRNAs, and 179 tRNAs.

**Table 1.** Read and assembly statistics for whole-genome sequencing of *Cosmosporella* sp. VM-42.

|  | Unfiltered | Filtered (> 2kb) |
| --- | --- | --- |
| No. of bases (Gbp) | 6.8 | 5.6 |
| No. of reads | 1,090,573 | 726,202 |
| N50 (kb) | 9.3 | 10.04 |
| Mean read length (bp) | 5,481.8 | 7,704.0 |
| Mean read quality | 17.6 | 19.2 |
| Coverage | 120 x |  |
| Polishing steps | Racon+Medaka |  |
| No. of contigs (>= 50000 bp) | 10 |  |
| Genome size (Mbp) | 39.07 |  |
| GC (%) | 48.11 |  |
| BUSCO (Ascomycota_odb10) (%) | 97.3 |  |
| Number of protein coding genes | 11,642 |  |
| Number of tRNA genes | 179 |  |
| Number of rRNA genes | 55 |  |
| Proteins with predicted Pfam domain | 9,842 |  |
| CAZymes | 415 |  |

Next, we performed multilocus phylogenetic analysis with the ITS, LSU, and *TUB2* loci of this fungus, using 23 other taxa in the *Nectriaceae* family, and *Fusarium oxysporum* Fo47 as an outgroup (Figure 1A). The resulting phylogenetic tree revealed that our fungal isolate fell into the same clade as *Cosmosporella,* a sister clade of various *Cosmospora* species. Furthermore, we observed that it clustered with one undefined species, ‘*Nectria*’ *flavoviridis* (IMI 338173), which is related to *Cosmosporella olivacea* in a strongly supported monophyletic lineage (Huang et al., 2018). Therefore, we further classified our fungal isolate as *Cosmosporella* sp. VM-42. Another undefined species, ‘*Cosmospora*’ *obscura* (MAFF 241484), clustered in the same clade indicating that this fungus with largely unresolved taxonomy might also be a *Cosmosporella* species.

**Figure 1.**
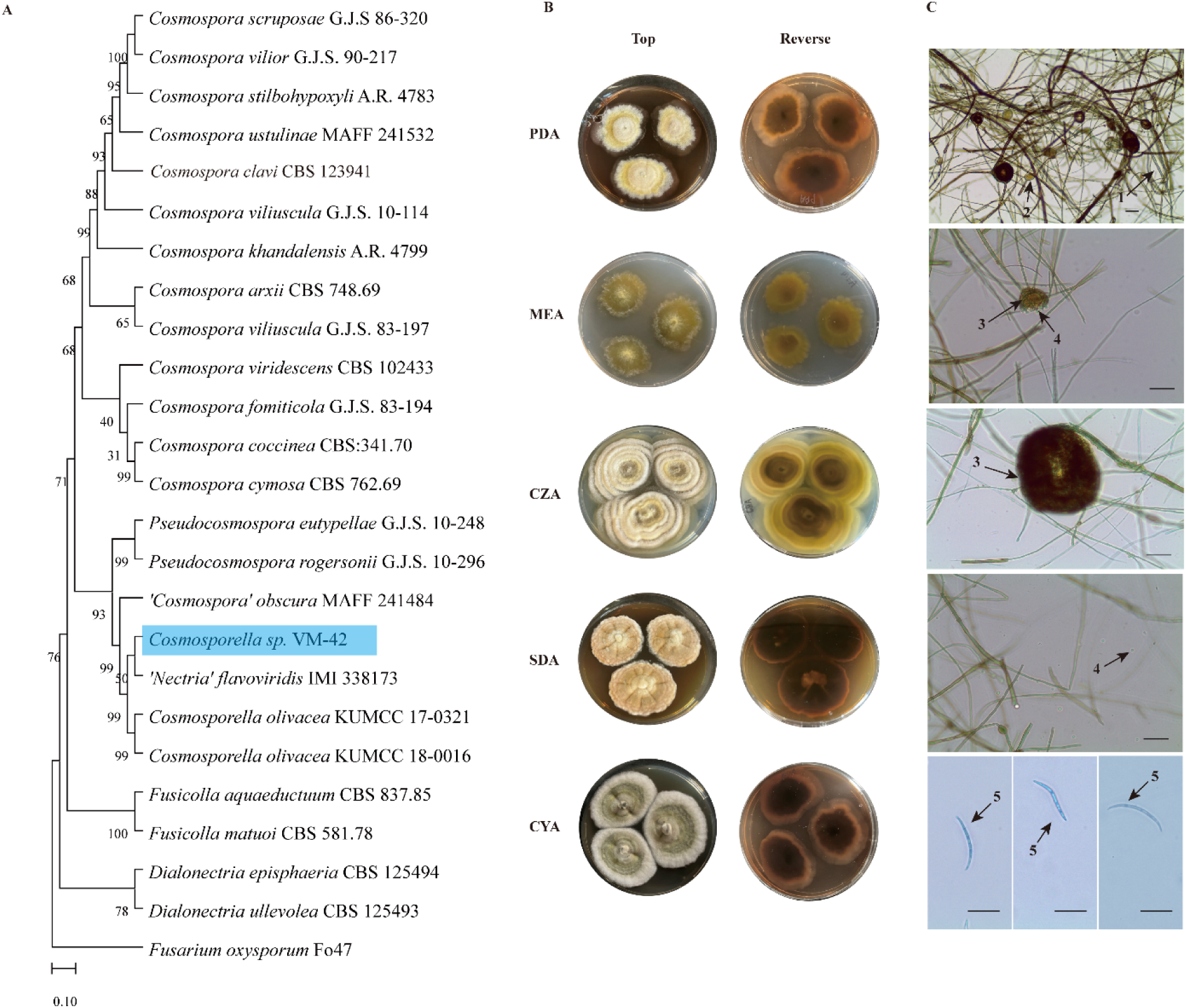
Multilocus Phylogeny and Morphological Analysis of *Cosmosporella* sp. VM-42. (A) Multilocus phylogeny analysis based on *tub2*, ITS, and LSU concatenated nucleotide sequences with *Fusarium oxysporum* Fo47 as outgroup. (B) Macroscopic and (C) microscopic characteristics of *Cosmosporella* sp. VM-42 upon growth on different culture media (PDA, MEA, CZA, SDA, and CYA) at 25°C for 28 days. (1) Hyphae; (2) Coiled hyphae; (3) Pycnidium; (4) Ovoid microconidia; (5) Macroconidia. Scale bars = 50 µm.

We further characterized *Cosmosporella* sp. VM-42 morphologically upon growth on different media (Figure 1B). Colonies grown on PDA plates reached a radius of 17 mm in 28 days at 25°C, with a white center and pale-yellow intervals, woolly texture, and white aerial hyphae. The surrounding medium became brown to maroon by diffusing pigments, and the reverse of the colonies showed a tan center fading into light red. Compared to other culture media, colonies grew slowest on MEA medium, attaining a radius of 14 mm after 28 days incubation. The center of the colony was smooth with poorly developed aerial mycelium around the edges. Both sides of the colony were yellow. Colonies on CZA plate grew faster than those on other media, reaching a radius of 22 mm in 28 days. They displayed apparent concentric rings with a greyish center and pale-yellow and white intervals, woolly texture and an aerial mycelium. The culture medium was not impregnated with pigments. The reverse of the colonies also displayed concentric rings with a brownish black center and yellow intervals. Colonies on SDA plates attained a radius of 17 mm in 28 days, slightly raised at the center, with orange-pink hyphae, a floccose texture and sulcate appearance. Their pigments were released into the culture medium, with a tan center and narrower light red edge in the reverse side. Colonies on CYA plates reached a radius of 19 mm, possessed an olivaceous-grey center surrounded by white mycelium, and were velvety textured with aerial mycelium. They displayed similar reverse morphology as the colonies on SDA and PDA media. Optical microscopy revealed the presence of pycnidia, ovoid microconidia, and macroconidia (Figure 1C).

### 3.2. Cosmosporella sp. VM-42 shows antibacterial activity against MSSA and MRSA strains

To investigate the antibacterial potential of *Cosmosporella* sp. VM-42, we grew the isolate in liquid culture and generated a crude EtOAC extract. This extract exhibited a strong inhibition effect on growth of *S. aureus* ATCC 29213 (92%). In contrast, we observed hardly any growth inhibition of *E. coli* ATCC 25922 (Figure 2A). Next, we tested the *Cosmosporella* sp. VM-42 extract against different MSSA and MRSA strains. The MIC of the crude EtOAc extract was 125 µg/mL for all tested MSSA strains and the MRSA strain USA300 (Figure 2B and Figure S2B). In comparison to MIC values reported in other studies (Arivudainambi et al., 2011; Phongpaichit et al., 2006), the crude extract of *Cosmosporella* sp. VM-42 showed intermediate-high antimicrobial activity against both MSSA and MRSA strains. Thus, we set out to investigate the secondary metabolites produced by this fungus to identify the bioactive compound(s) responsible for the antibacterial activity.

**Figure 2.**
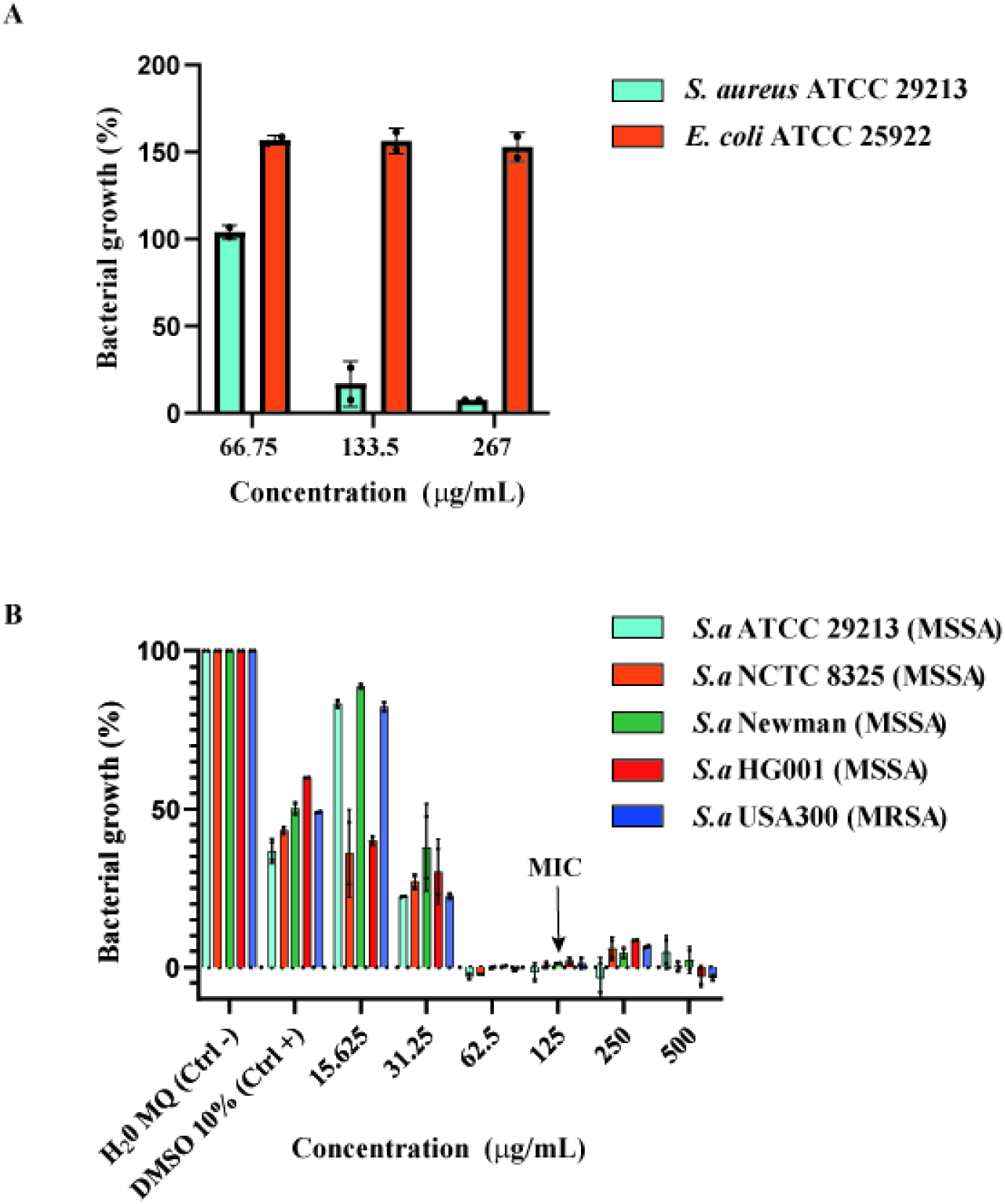
Antibacterial susceptibility testing of *Cosmosporella* sp. VM-42. (**A**) Selective inhibitory activity of the sEtOAc fraction of *Cosmosporella* sp. VM-42 against *S. aureus* ATCC 29213; (**B**) Dose-dependent inhibition of the sEtOAc fraction against MSSA and MRSA strains quantified as % bacterial growth based on OD_600_ (MIC: 125 µg/mL); bars represent mean ± SD of technical duplicates.

### 3.3. Cosmosporella sp. VM-42 produces multiple secondary metabolites

To activate the expression of potentially silent biosynthetic gene clusters (BGCs) and obtain more secondary metabolites, we exposed *Cosmosporella* sp. VM-42 to two frequently used epigenetic modulators, sodium butyrate (SB) and procaine hydrochloride (PH). After 14 days of cultivation at 25℃ in presence of these modulators, we analyzed the phenotypes and metabolomes of the colonies (Figure S3). LR-LC-MS of three biological samples for each condition showed similar peak retention times and mass-to-charge (*m/z*). Before the addition of chemical modifiers, *Cosmosporella* sp. VM-42 produced several metabolites visible as distinct peaks in the total ion chromatograms (TICs, Figure S3), including dominant peaks with *m/z* 195.17 (rt 13.50 min) and *m/z* 277.31 (rt 13.77 min), as well as some relatively small peaks with *m/z* 181.06 (rt 10.59 min), *m/z* 515.10 (rt 12.48 min), *m/z* 277.23 (rt 13.97 min), *m/z* 535.27 (rt 16.39 min), *m/z* 517.10 (rt 19.14 min), *m/z* 519.17 (rt 19.38 min), *m/z* 882.57 (rt 21.33 min), *m/z* 279.07 (rt 21.50 min), *m/z* 874.67 (rt 21.62 min), and *m/z* 888.63 (rt 22.00 min). With increasing concentrations of SB, the fungus produced more black and brown pigments, especially at 10 mM SB, forming distinct black droplets. However, the higher concentration of 100 mM SB did not elicit more pigment production and instead inhibited growth and pigment production and release. LR-LC-MS analysis revealed that SB at 10 mM did not only enhance the intensity of the afore-mentioned peaks in the PDA sample group, but also triggered the production of new peaks, especially two prominent peaks with *m/z* 293.18 (rt 11.21 min) and *m/z* 295.19 (rt 18.71) (Figure S3A). With increasing concentration of the second epigenetic modifier, PH, the fungal mycelium turned from yellow to grey. Again, the highest concentration, 100 mM, completely inhibited fungal growth. Based on the TICs, 10 mM PH increased the production of several main compounds already present in the PDA control group and elicited three dominant changes, namely the appearance of two broad peaks with *m/z* 293.03 (rt 12.40 min - 13.00 min) and *m/z* 277.16 (rt 19.40 min - 20.40 min), one peak with *m/z* 632.23 (rt 13.23), and one sharp peak with *m/z* 834.42 (rt 16.11 min) (Figure S3B).

To identify the chemical diversity of secondary metabolites in our *Cosmosporella* sp. VM-42 isolate, we also collected HR-LC-MS/MS data of EtOAc extracts of the fungus after treatment with 10 mM SB and PH, and conducted a Global Natural Products Social Molecular Network analysis (GNPS-MN) followed by compound identification and prediction with SNAP-MS and SIRIUS 5.8.6. Nodes present in the uninoculated PDA control groups with 10 mM SB and PH were omitted from the final molecular network with 874 nodes (Figure 3). Based on the combined chemical information from GNPS-MN, SNAP-MS and SIRIUS, as well as neighborhood inference, we putatively identified 26 compounds in the network (Table S3, Supplementary Note S1). For instance, in cluster 1, two nodes with *m/z* 888.547 and *m/z* 874.531 are predicted by GNPS to be [M + H]^+^ adducts of _130100 (**1**) and _130099 (**2**), also known as W493B and W493A. Node *m/z* 846.501 (**3**) shows -C_3_H_6_ and -C_2_H_4_ differences with **1** and **2**. Although most of its MS2 fragments are identical to those of **1** and **2**, the existence of the *m/z* 435.1892 and *m/z* 253.1192 fragments indicates that L-Ile in **1** and L-Val in **2** were replaced by L-Ala in **3**, a putative novel lipopeptide. Node *m/z* 886.532 (**4**) has one -H_2_ group less than **1**, indicating the addition of one conjugated double bond, and it is predicted to be W493D according to SNAP-MS analysis (Lv et al., 2015). Node *m/z* 904.543 (**5**) contains one more oxygen atom than **1**, and MS2 fragments with *m/z* 493.2313 and *m/z* 311.1610 indicate the addition of a hydroxyl group on the benzene ring, and it is predicted to be Acuminatum F (Zhong et al., 2023). Node *m/z* 860.517 (**6**) has less -C_2_H_4_ and -C_2_H_2_ groups than **1** and **4**, respectively. Based on the fragments with *m/z* 586.4156, *m/z* 572.3724, *m/z* 548.3433, *m/z* 438.2981, *m*/*z* 420.2863, *m*/*z* 367.2600, *m*/z 349.2497, *m/z* 296.2228, and *m/z* 278.2125, a 3-hydroxy-4-methyldodecanoic acid moiety replaces the original fatty acid side chain, suggesting that it represents W493C (**6**) (Lv et al., 2015). The node with *m/z* 890.527 (**7**) has one more oxygen atom than **2,** one less -CH_2_ group than **5**, and it has an indirect relationship with **1**, indicating that it consists of L-Val and 3, 4-dihydroxy-phenylalanine units instead of L-Ile and 4-hydroxy-phenylalanine moieties. Its MS2 fragments, including fragments with *m/z* 479.2173, *m/z* 407.4955, *m/z* 377.2819, *m/z* 324.2549, and *m/z* 297.1458, provide evidence to support this view. Therefore, compound **7** is predicted to be 3-((3*S*,6*R*,9*S*,12*R*,15*S*,18*R*,22*S*)-6-(3,4-dihydroxybenzyl)-22-((*R*)-dodecan-2-yl)-18-((*R*)-1-hydroxyethyl)-3-isopropyl-12,15-dimethyl-2,5,8,11,14,17,20-heptaoxo-1-oxa-4,7,10,13,16,19-hexaazacyclodocosan-9-yl)propanamide, a putative new compound. Similarly, node *m/z* 872.516 (**8**) shows one -H_2_ group difference compared to **2**, and fragments with *m/z* 574.3671, *m/z* 482.3228, *m/z* 464.3130, *m/z* 446.3032, *m/z* 393.2762, *m/z* 322.2386, and *m/z* 304.2280 indicate the existence of one double bond in the fatty acid chain. Node *m/z* 804.454 (**9**) has one -C_5_H_10_ group less than **2**, and fragments with *m/z* 506.2964, *m/z* 414.2623, *m/z* 396.2505, *m/z* 378.2410, *m/z* 343.2241, *m/z* 325.2138, *m/z* 307.2030, *m/z* 254.1763, *m/z* 236.1658, and *m/z* 118.0868, indicating the existence of a 3-hydroxy-4-methyl-nonacanoic acid moiety in **9**. Node *m/z* 846.5 (**10**) has one -C_2_H_4_ group difference compared to **2**, and a 3-hydroxy-4-methyldodecanoic acid moiety highly likely exists as indicated by the fragments with *m/z* 666.3894, *m/z* 584.3693, *m/z* 572.3988, *m/z* 548.3491, *m/z* 438.2972, *m/z* 420.2876, *m/z* 391.1996, *m/z* 367.2603, *m/z* 349.2497, *m/z* 296.2229, and *m/z* 278.2125. They are putative new compounds. Detailed annotations of other compounds are described in Supplementary Note S1. Structures of the predicted and inferred secondary metabolites and their MS2 fragments are displayed in Figure 4.

**Figure 3.**
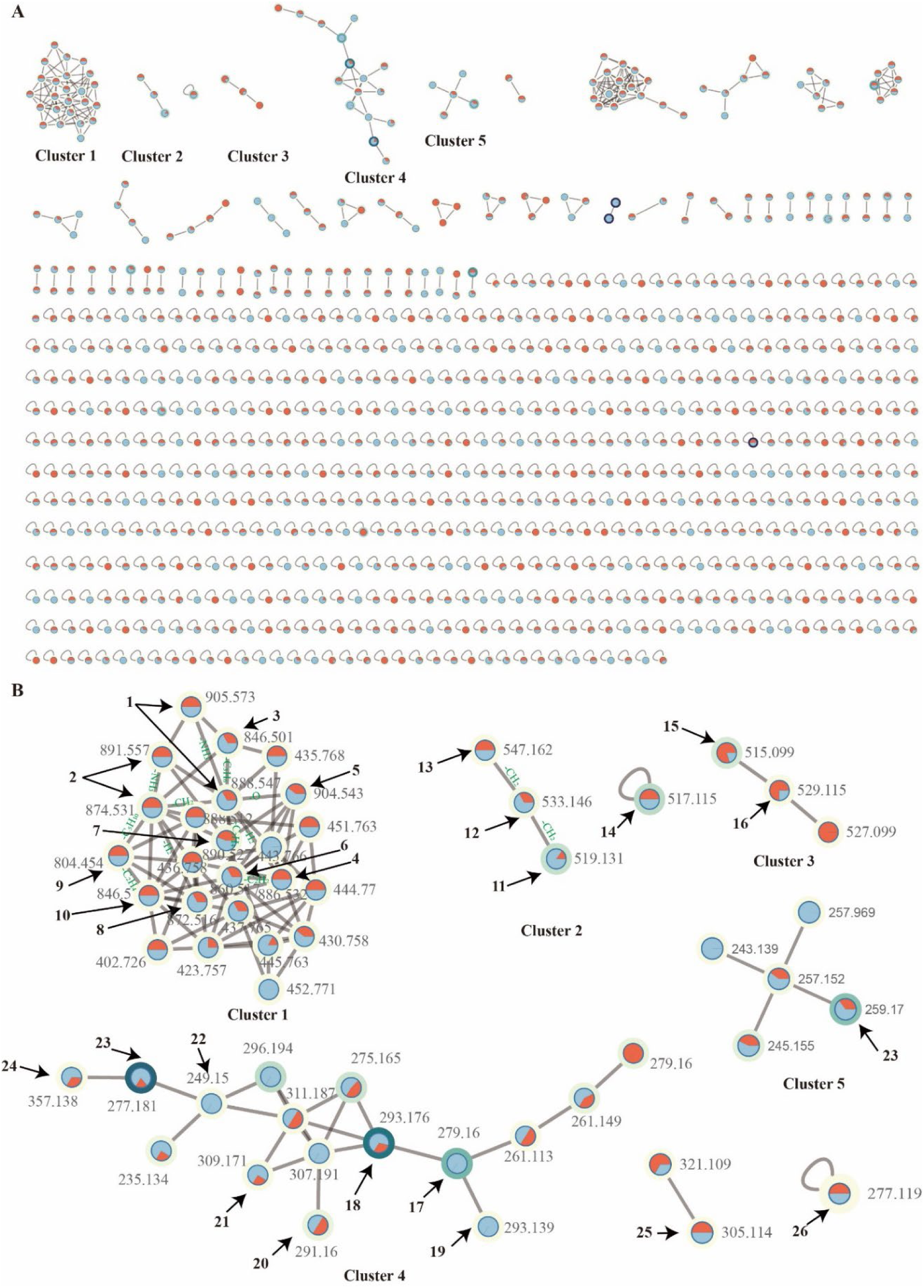
Molecular network of *Cosmosporella* sp. VM-42 secondary metabolites before and after the treatment with 10 mM of the small molecule epigenetic modifiers sodium butyrate (SB) or procaine hydrochloride (PH). (A) Overview of the molecular network; (B) Zoom in on clusters 1-5 with nodes putatively matched to compound structures (see Figure 4) based on GNPS, SNAP-MS, and SIRIUS analysis. Node labels indicate the *m/z* values in MS1, node colours indicate the presence of the feature in the extracts of fungus treated with PH (blue) or SB (red). Border paint, ranging from yellow to purple, indicates an increase in the precursor abundance in samples based on peak area intensity. Edges represent the structural similarity between nodes.

**Figure 4.**
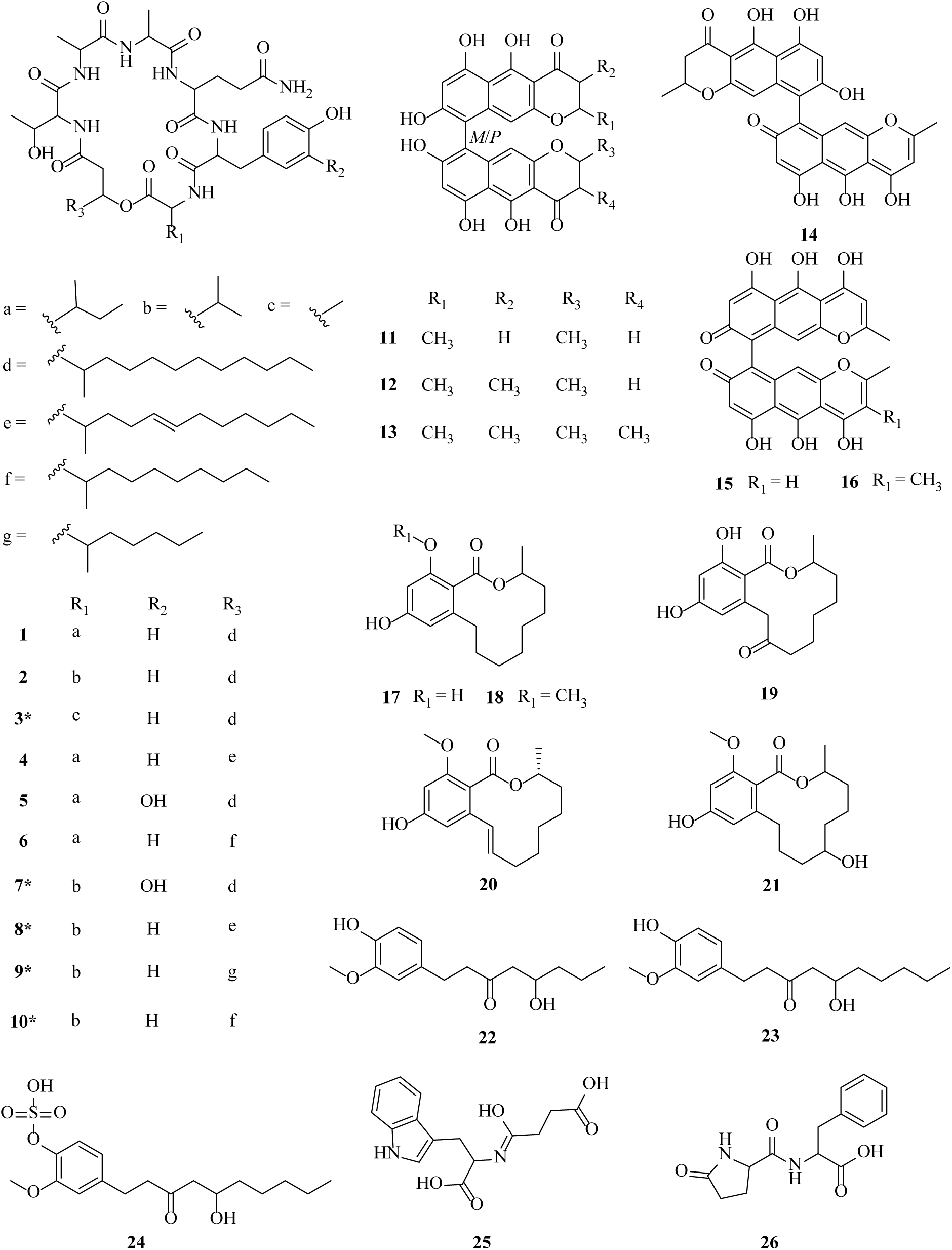
Chemical structures of secondary metabolites putatively identified in *Cosmosporella* sp. VM-42 extracts after treating the fungus with sodium butyrate or procaine hydrochloride (* indicates putative novel compounds).

Despite our compound discovery efforts, some peaks with high intensities in the TIC remained unidentified, especially the peaks with *m/z* 195.1019 (rt 14.83 min) and *m/z* 279.1959 (rt 20.49 min) detected before epigenetic manipulation, the newly produced peak with *m/z* 295.1909 (rt 18.98 min) upon exposure to 10 mM SB, and the enhanced peaks with *m/z* 632.2374 (rt 10.75 min), *m/z* 834.4155 (rt 13.83 min), *m/z* 277.1808 (rt 11.47 min), and *m/z* 293.1756 (rt 11.48) observed in the group exposed to 10 mM PH. These compounds are promising candidates for the discovery of novel secondary metabolites by purification and structure elucidation. For instance, the peaks with *m/z* 834.4155 and *m/z* 632.2374 are predicted to be peptides by SIRIUS, but no corresponding candidate compound has been found in the database, indicating that they are highly likely to be novel compounds.

### 3.4. Isolation, structural identification and antibacterial effect of compound 27

Since the peak with *m/z* 195.1026 was dominant in the crude extract of our *Cosmosporella* sp. VM-42 isolate and preliminary tests showed that fractions containing this peak possessed a strong inhibitory activity towards *B. subtilis*, we decided to isolate this compound. From 2 L of culture media, we purified compound **27** as a brown green powder, with UV absorption maxima at 225.63 and 330.63 nm and *m/z* 195.1026 ([M + H]^+^ adduct; chemical formula C_11_H_14_O_3_, 194.0948). The ^1^H, ^13^C, and 2D NMR data for compound **27** are shown in Table 2 and Figure S4-S8. Although the carbon spectrum signal was weak, we still found relevant 2D NMR signals and assigned them. Based on literature (Reveglia et al., 2021), we identified compound **27** as nectriapyrone, which was previously isolated from the fungus *Diaporthe eres*. The chemical structure and key 2D NMR correlations of compound **27** are displayed in Figure 5.

**Figure 5.**
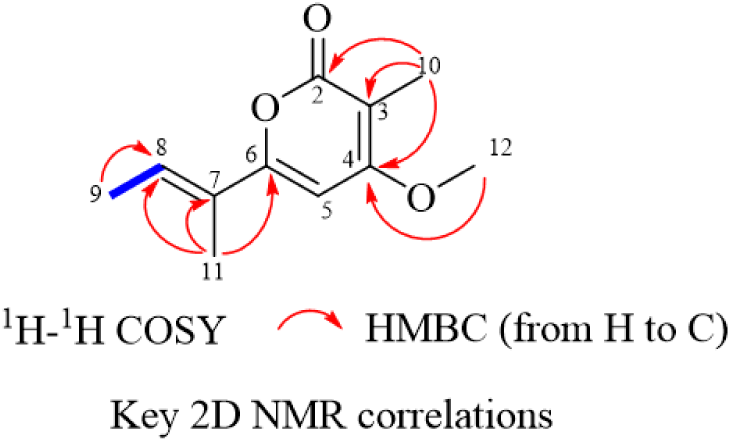
Chemical structure and key 2D NMR correlations of compound 27.

**Table 2.** ^1^H NMR (600 MHz), ^13^C NMR (150 MHz), ^1^H-^1^H COSY, and HMBC data of compound 27 (in DMSO-d6).

| No. | $\delta_{\text{C}}$, type | $\delta_{\text{H}}$ (mult., $J$ in Hz) | HMBC | $^1\text{H}$-$^1\text{H}$ COSY |
| --- | --- | --- | --- | --- |
| 2 | 164.0, C | - | Me-10 | - |
| 3 | 100.11, C | - | Me-10 | - |
| 4 | 166.6, C | - | Me-10, OMe-12 | - |
| 5 | 100.22, CH | 6.48, s | - | - |
| 6 | 159.68, C | - | Me-11 | - |
| 7 | 128.28, C | - | Me-11 | - |
| 8 | 128.5, CH | 6.50, m | Me-9, Me-11 | Me-9 |
| 9 | 14.29, $\text{CH}_3$ | 1.83, d (7.1) | H8 | H8 |
| 10 | 8.79, $\text{CH}_3$ | 1.78, s | C2, C3, C4 | - |
| 11 | 12.33, $\text{CH}_3$ | 1.89, s | C6, C7, C8 | - |
| 12 | 57.18, $\text{OCH}_3$ | 3.94, s | C4 | - |

- not detected.

Next, we determined the MIC and MBC of compound **27** against the MRSA strains USA 300, D15-CA, D17-HA and MSSA ATCC 29213 and observed that it ranges from 125 to 62.5 µg/mL (Table 3). These results show that compound **27** has an *in vitro* antibacterial inhibitory effect against the tested MRSA and MSSA strains in a dose-dependent manner (Figure 6A), contributing to the antibacterial properties of the crude EtOAC extract from *Cosmosporella* sp. VM-42.

**Figure 6.**
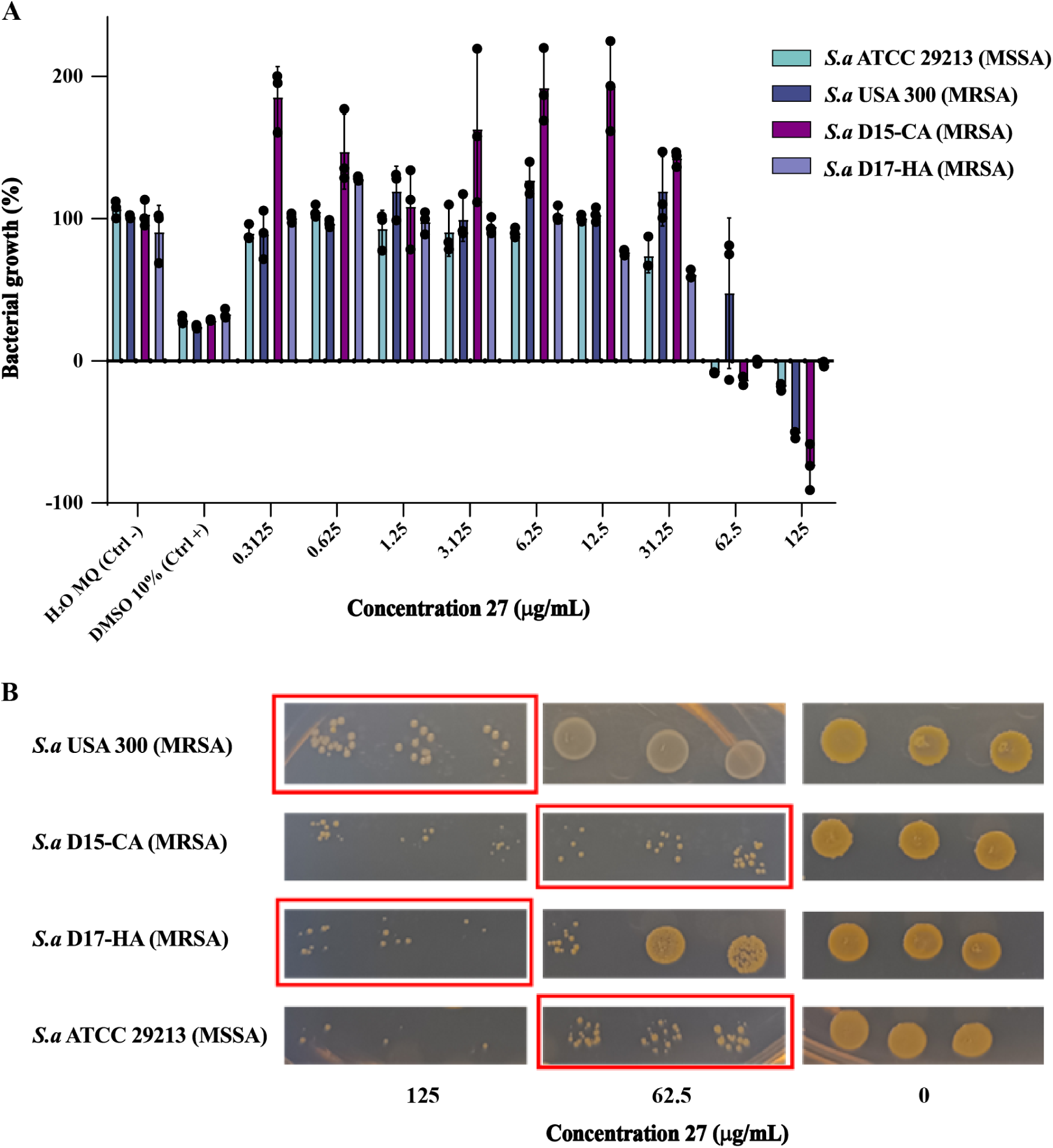
Antibacterial effect of compound 27 produced by *Cosmosporella* sp. VM-42 against MSSA and MRSA strains. (A) Dose-dependant inhibitory effect of compound 27 with MIC shown next to each strain. Bars represent mean +/-SD, n=3. (B) Bactericidal effect of compound **27**. MBC values are marked within red boxes.

**Table 3.**
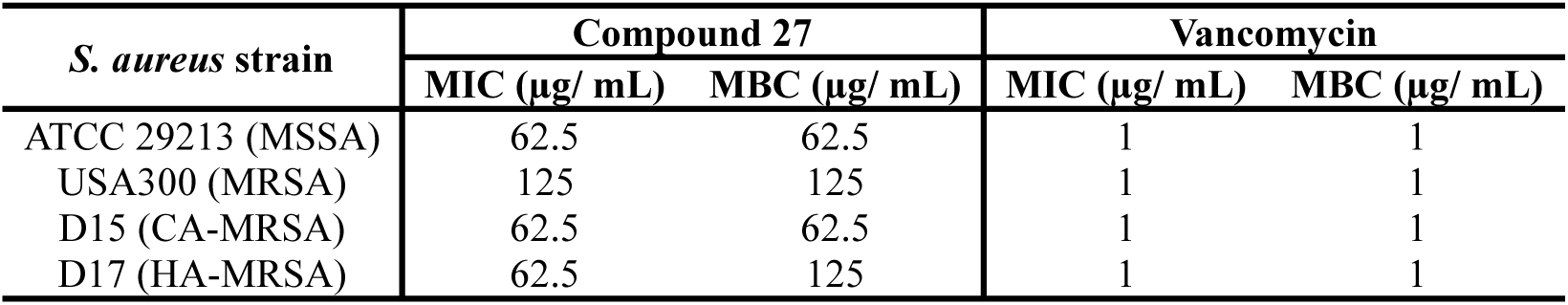
MIC and MBC values of compound 27 against MSSA and MRSA strains.

| *S. aureus* strain | Compound **27** |  | Vancomycin |  |
| --- | --- | --- | --- | --- |
| | MIC ($\mu\text{g}/\text{mL}$) | MBC ($\mu\text{g}/\text{mL}$) | MIC ($\mu\text{g}/\text{mL}$) | MBC ($\mu\text{g}/\text{mL}$) |
| ATCC 29213 (MSSA) | 62.5 | 62.5 | 1 | 1 |
| USA300 (MRSA) | 125 | 125 | 1 | 1 |
| D15 (CA-MRSA) | 62.5 | 62.5 | 1 | 1 |
| D17 (HA-MRSA) | 62.5 | 125 | 1 | 1 |

Importantly, the results presented in Figure 6B imply that compound **27** exerts a bactericidal activity against the tested MSSA and MRSA strains. Nonetheless, it should be noted that the antibacterial potency of compound **27** against MSSA and MRSA strains is relatively low, which is consistent with previous reports on the antibiotic activity of nectriapyrone against *S. aureus* at a concentration of 30 ppm (M.S.R. Nair and Susan T. Carey, 1975). This also indicates that other antibiotic compounds in the crude extract of *Cosmosporella* sp. VM-42 probably contribute to the potency we observed in our initial experiments with the crude EtOAC extract.

### 3.5. The genome of Cosmosporella sp. VM-42 encodes multiple enzymes of biotechnological interest

To evaluate the metabolic potential of *Cosmosporella* sp. VM-42, we further analyzed the whole-genome sequence of this fungus. First, we first performed a functional annotation of the predicted genes for the main gene ontologies (Supplementary dataset 1, Figure S9A) and assembled the mitochondrial genome (Figure S9B). Second, we searched the genome for BGCs using AntiSMASH. In total, 35 BGCs were predicted and their putative products were classified based on their core biosynthetic genes (Table S4). Among them, 11 BGCs display high similarities with reported BGCs in the MiBIG database (Terlouw et al., 2023), and the chemical structures of the products of these known clusters are shown in Figure S10. In the following, we address those BGCs that could be associated with the compounds in the major clusters of our molecular network.

Region 11.4 is a hybrid non-ribosomal peptide synthetase (NRPS)/ Type1 polyketide synthase (T1PKS)-encoding BGC. KnownClusterBlast showed that 100% of the genes in BGC0002188 from *Fusarium pseudograminearum CS3096*, which is responsible for producing the lipopeptides W493B and W493A, has a significant match with this query sequence (Figure 7A). The NRPS and PKS in both clusters show 71% and 84% amino acid identities, respectively. The biosynthetic pathway of W493A and W493B as displayed in Figure 8A was previously proposed based on mutagenesis studies, transcript and metabolite analysis (Sørensen et al., 2014), and heterologous expression of the intact BGC (Nielsen et al., 2019). It is plausible that region 11.4 gives rise to the same type of compounds as BGC0002188. The presence of additional genes with predicted functions, including short-chain reductase and triglyceride lipase activity, however, suggests that other derivatives of these lipopeptides could be produced. This is in good agreement with our molecular networking analysis, where an entire cluster of related compounds was identified (cluster 1, compounds **1**-**10**).

**Figure 7.**
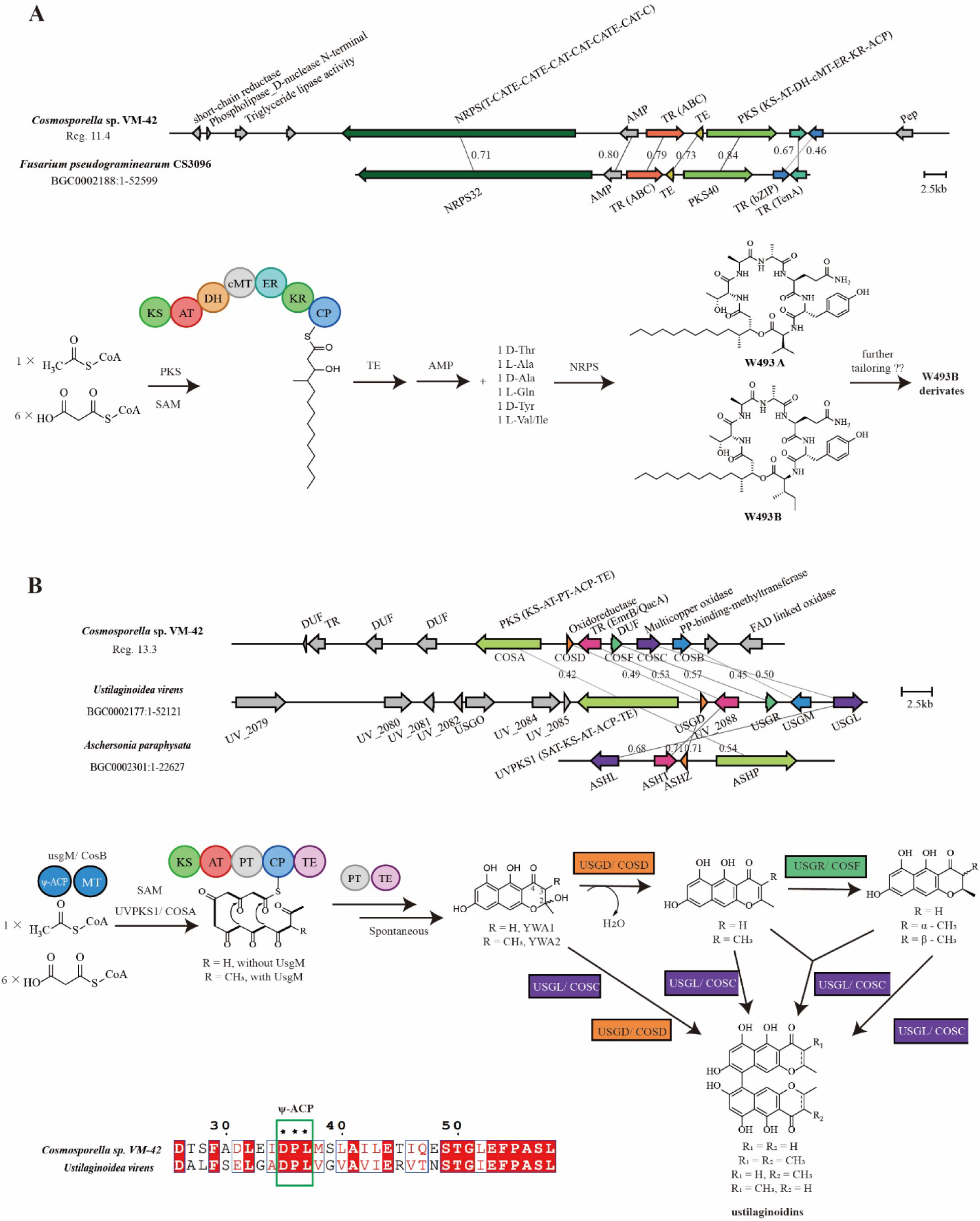
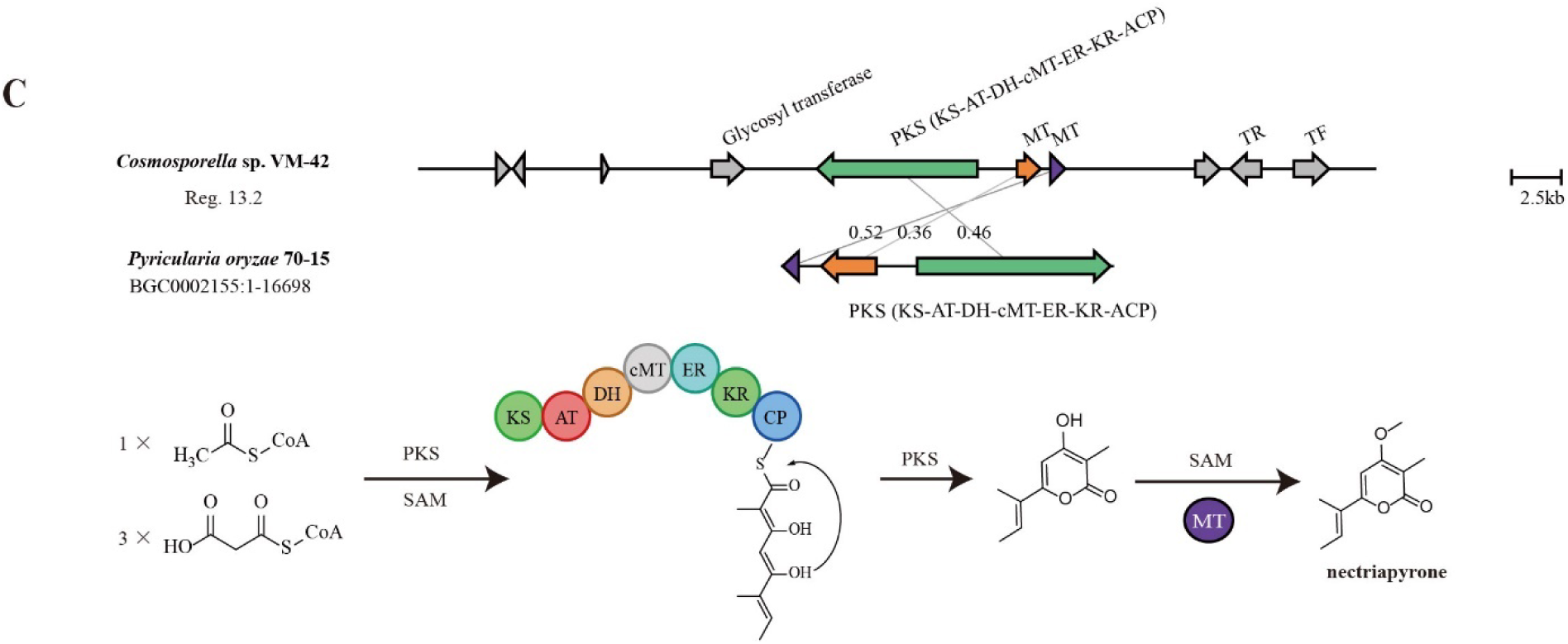
BGCs in the *Cosmosporella* sp. VM-42 genome compared to homologous BGCs and their proposed products. (A) Region 11.4 is homologous to a BGC from *Fusarium pseudograminearum* CS3096; (B) Region 13.3 is homologous to BGCs from *Ustilaginoidea virens* and *Aschersonia paraphysata*; (C) Region 13.2 is homologous to a BGC from *Pyricularia oryzae* 70-15. Gene cluster comparison and visualization were performed by clinker on the CAGECAT web server (https://cagecat.bioinformatics.nl/). Genes are represented as arrows with the predicted gene product annotated above or below. The small numbers on the lines connecting genes are amino acid sequence identities expressed in decimal fractions.

Region 13.3 encodes a non-reducing (nr)PKS. KnownClusterBlast revealed that 46% of the genes in an ustilaginoidin N BGC (BGC0002177) from *Ustilaginoidea virens* and 100% of the genes in a (*P*)-ustilaginoidin A BGC (BGC0002301) from *Aschersonia paraphysata* have significant matches to the query sequence. Ustilaginoidins are mycotoxins that feature a dimeric naphtho-γ-pyrone skeleton (Xu et al., 2021). The core PKS gene encoded in region 13.3 shares 42% amino acid sequence identity with UvPKS1 from *Ustilaginoidea virens*. According to the literature (Y. Li et al., 2019; Obermaier et al., 2019; Xu et al., 2021), this PKS produces an Acyl Carrier Protein (ACP)-bound heptaketide, which is further processed by the product template (PT) and thioesterase (TE) domains before being released as a phenolic ketone that spontaneously ketalizes to the monomeric naphtho-γ-pyrone YWA1. A methyltransferase (MT) with a phosphopantetheine binding site in region 13.3 shows 45% amino acid identity with UsgM from *Ustilaginoidea virens*. Based on sequence alignments using CLUSTALW and ESPript 3, this protein has a similar ψ-ACP-MT didomain architecture as UsgM, including a D-P-L motif in the phosphopantetheine modification site instead of the canonical D-S-L motif (Figure 7B). The ψ-ACP domain is important for forming the 3-methyl derivative YWA2, which is also an intermediate in tricholignan A biosynthesis (Chen et al., 2019; Xu et al., 2021). The enzymes encoded by *cosD*, *cosF*, and *cosC* in region 13.3 share 49%, 57%, and 50% amino acid identities with the enzymes encoded by *usgD*, *usgR*, and *usgL*, respectively. Gene-deletion and heterologous expression in *Aspergillus oryzae NSAR1* (Xu et al., 2021) elucidated the roles of these enzymes in dimeric naphtho-γ-pyrone biosynthesis: UsgD is a dehydratase enzyme accelerating the formation of Δ^2^-derivates, which can also form slowly and spontaneously (Xu et al., 2021). UsgR is an ene-reductase required for the asymmetric reduction of the 2, 3-double bond to yield epimers at C-3 position (Xu et al., 2021). UsgL is a laccase responsible for the dimerization of monomers and is known for its remarkable flexibility in producing two atropisomers: at lower enzyme concentration *P*-ustilatinoidin A is the preferred product and at higher concentrations its *M*-atropisomer (Obermaier et al., 2019). In addition, *in vitro* experiments revealed that the atroposelective dimerization of monomers by the laccase UsgL preferred 3-nonmethyl-over 3-methyl-YWA1, 2,3-saturated over 2,3-unsaturated YWA1, YWA1 over its dehydrated counterpart, and the dehydrated derivative of 3-methyl YWA1 over 3-methyl YWA1 (Xu et al., 2021). Secondary metabolite analysis in *Cosmosporella* sp. VM-42 indicated the existence of 4 other types of congeners derived from ustilaginoidin A by Δ^2^ double-bond saturation, methylation at position 3, or a combination of these two characteristics. Although we did not detect all naphtho-γ-pyrones that are known to be formed in *Ustilaginoidea virens*, the existence of similar biosynthetic genes in the *Cosmosporella* sp. VM-42 genome indicates the potential for producing a variety of dimeric naphtho-γ-pyrones, as we found in clusters 2 and 3 of the molecular network (compounds **11**-**16**).

The BGC in region 13.2 includes a core gene encoding a highly-reducing (hr)PKS, which shares 46% identity with the PKS from *Pyricularia oryzae* 70-15 involved in the biosynthesis of (*E*)-6-(but-2-en-2-yl)-4-hydroxy-3-methyl-2*H*-pyran-2-one by consuming 1 acetyl-CoA and 3 malonyl-CoA (Figure 7C). Two O-methyltransferases are also encoded in this BGC, and one of them shares 52% identity with NEC2 in *P. oryzae* 70-15, which is responsible for further methylation at position 4 to form nectriapyrone (compound **27**), which can inhibit the growth of *Streptomyces griseus* (Motoyama et al., 2019). Nectriapyrone has been found to be converted to nectriapyrone D or gulypyrone B in the presence of a hydroxylase in *P. oryzae* 70-15 (Motoyama et al., 2019). Since we did not detect such hydroxylase-encoding genes in the BGC of *Cosmosporella* sp. VM-42, we assume that hydroxylated derivatives are likely not formed in our fungus, which is also in agreement with the absence of such derivatives in the molecular network.

In addition to the BGCs mentioned above, *Cosmosporella* sp. VM-42 still has enormous potential for the synthesis of polyketides and peptides, which we describe in more detail in Supplementary Note S2 (Figure S11). Furthermore, some gene clusters with unknown function also attracted our attention, since they could give rise to compounds putatively identified in the metabolome of *Cosmosporella* sp. VM-42. For instance, the new peaks of *m/z* 834.4155 and *m/z* 632.2374 arising upon treatment with 10 mM procaine hydrocholoride are predicted to be novel peptides. Considering the biosynthetic logic, these compounds could be related to the NRP/metallophore hybrid BGC in region 16.2 or the NRP BGC in region 16.3. The expression of these BGCs might be influenced by procaine hydrocholoride, but these assumptions require further verification by compound isolation and transcriptomic analysis.

## 4. Discussion

Our present genomic and metabolomic analyses show that *Cosmosporella* sp. VM-42 has the potential to produce a diverse set of interesting compounds and enzymes of biotechnological interest. Here, we want to highlight several striking secondary metabolites and BGCs in this fungus.

Compounds **1**-**10** in this study belong to the cyclodepsipeptides, among which five were identified as putative novel compounds. The BGC in region 11.4 is predicted to be responsible for producing this class of compounds. Compounds **1**, **2**, **4**, **5**, **6** were identified as W493B, W493A, W493D, Acuminatum F, and W493C, respectively. These compounds have been reported to exhibit antifungal activities (Zhong et al., 2023), which generates considerable interest due to the emergence of fungal pathogens and the high rate of antifungal resistance. Therefore, in further studies it would be highly interesting to explore the antifungal activities of the five putative novel cyclodepsipeptides (**3**, **7**, **8**, **9**, and **10**). Moreover, several studies have reported the anti-MRSA activities of cyclodepsipeptides (Bionda et al., 2013; Kadouri and Shanks, 2013). This suggests that the cyclodepsipeptides identified in our study may contribute to the strong antibacterial effect of the crude *Cosmosporella* sp. VM-42 extract and may serve as potential lead compounds with antifungal and antibacterial properties.

*Cosmosporella* sp. VM-42 also produced ustilaginoidins, a class of bis-naphtho-γ-pyrone mycotoxins (compounds **11**-**16**), and the BGC in region 13.3 is likely responsible for their biosynthesis. Compounds of this class occur widely in fungi and they exhibit a variety of bioactivities, such as cytotoxicity, phytotoxicity, inhibition of the HIV-1 integrase, antitumor and antibacterial activity (Lu et al., 2015; Singh et al., 2003; Ugaki et al., 2012). Notably, specific ustilaginoidins, such as ustilaginoidins D, E, and G, are reported to potent antibacterial activity (Lai et al., 2019). These findings underscore the potential of ustilaginoidins for applications in medicine and agriculture.

Compounds **17**-**21** are lasiodiplodins, belonging to the family of macrolides, which also possess various biological activities (Kuttikrishnan et al., 2022; Luo et al., 2023; Zhang et al., 2017) and one lasiodiplodin was reported to exhibit anti-MRSA with MIC 128 mg mL^−1^ (Rukachaisirikul et al., 2009). The biosynthesis of lasiodiplodins is still unresolved, but it is predicted to involve similar steps as the biosynthesis of 14-membered macrolides, such as zearalenone (Kim et al., 2005; Lysøe et al., 2006) and hypothemycin (Reeves et al., 2008). However, the corresponding genes and enzymes for lasiodiplodin biosynthesis have not yet been identified and studied. At the beginning of the proposed pathway (Kashima et al., 2009), five intact acetyl-CoA molecules are condensed by a hrPKS to yield a pentaketide acyl intermediate. The acyl chain is transferred to an nrPKS catalyzing three condensations with malonyl-CoA and an aldol condensation yielding a resorcylic acyl intermediate. This intermediate undergoes intramolecular cyclization by nrPKS, leading to the formation of lasiodiplodin. Region 14.2 encodes the only nrPKS in the *Cosmosporella* sp. VM-42 genome, but there is no evidence that an hrPKS is also encoded in this BGC. We therefore hypothesize that one of the hrPKSs encoded in regions 9.6 or 9.8 could work together with the nrPKS encoded in region 14.2 for the biosynthesis of lasiodiplodins.

Lastly, we purified the most abundant peak present in the crude extract of our *Cosmosporella* sp. VM-42 isolate (*m/z* of 195, rt 14.83 min), and identified it as nectriapyrone (compound **27**). Region 13.2 of *Cosmosporella* sp. VM-42 is predicted to produce nectriapyrone with 100% synteny compared to the MiBIG entry. Nectriapyrone has a broad spectrum of biological activities. It showed relevant cytotoxic activity against both human T leukemia and melanoma tumor cell lines, suggesting the potential use of this compound as a hit for the development of new anticancer drugs (Guimarães et al., 2008). Furthermore, nectriapyrone showed selective inhibition of butyrylcholinesterase, which is an important drug target for the treatment of Alzheimer’s disease (dos Santos et al., 2022). In our present study, it shows anti-MSSA and -MRSA activity via an unknown bactericidal mechanism. Currently, our understanding of the biological activities of nectriapyrone is still very limited, making it an interesting candidate for further investigation.

## 5. Conclusions

We isolated an endophytic fungus from *Vinca minor* and evaluated the antibacterial effects of its EtOAc extract, showing strong inhibition of MSSA and MRSA strains. We further obtained a high-quality whole-genome sequence of this endophytic fungus and classified it as *Cosmosporella* sp. VM-42. We identified 35 BGCs in the genome, some of which were predicted to encode known compounds, such as W493A, W493B, ustilaginoidin N, and nectriapyrone. These findings indicate that this fungus possesses great potential to produce multiple antimicrobial compounds.

We profiled the secondary metabolites of *Cosmosporella* sp. VM-42 by an untargeted metabolomic and molecular network analysis. Our results reveal that this fungus could produce a wide variety of bioactive compounds. Altogether, we putatively identified 26 compounds, including cyclodepsipeptides, ustilaginoidins, and lasiodiplodins, many of which are known to be antimicrobial compounds. We purified and characterized the most abundant compound from a crude extract of *Cosmosporella* sp. VM-42 and identified it as nectriapyrone (compound **27**), which shows anti-MSSA and -MRSA activity via an unknown bactericidal mechanism.

Overall, we systematically explored the *Cosmosporella* sp. VM-42, a thus far underexplored species, revealing its great potential for antimicrobial and biosynthetic applications. We anticipate that our study will facilitate future research on novel compounds and applications from endophytic fungi, such as *Cosmosporella* sp.. To open up possibilities for new drug discovery further targeted genome mining will be necessary, involving gene knockout studies, heterologous expression of microbial genes, regulation of promoters, and induction of mutations to biosynthesize new bioactive compounds.

## Supplementary Materials

Supplementary Information file with Table S1. List of species in *Nectriaceae* from NCBI used for phylogenetic analysis; Table S2. Quality metrics for isolated genomic DNA of *Cosmosporella sp.* VM-42; Table S3. Molecular networking-based identification of secondary metabolites in *Cosmosporella* sp. VM-42; Table S4. Biosynthetic gene clusters of *Cosmosporella* sp. VM-42 predicted by antiSMASH version 7.1.0; Figure S1. Contigs of *Cosmosporella* sp. VM-42 visualized by Bandage; Figure S2. Antimicrobial activity of *Cosmosporella* sp. VM-42; Figure S3. *Cosmosporella* sp. VM-42 with sodium butyrate (SB; A) and procaine hydrochloride (PH; B); Figure S4. ^1^H NMR spectrum of compound **27** in DMSO-d6 (600 MHz); Figure S5. ^13^C NMR spectrum of compound **27** in DMSO-d6 (150 MHz); Figure S6. ^1^H-^1^H COSY spectrum of compound **27** in DMSO-d6; Figure S7. HSQC spectrum of compound **27** in DMSO-d6; Figure S8. HMBC spectrum of compound **27** in DMSO-d6; Figure S9. Functional genome annotation of *Cosmosporella* sp. VM-42; Figure S10. Compounds produced by BGCs that were returned as most similar entries in MiBIG for certain BGCs detected in the *Cosmosporella* sp. VM-42 genome based on antiSMASH analysis; Figure S11. BGCs in the *Cosmosporella* sp. VM-42 genome compared to homologous BGCs and their proposed products; Note S1. Detailed annotations of putatively identified other compounds in the molecular network; Note S2. Other biosynthetic potential of *Cosmosporella* sp. VM-42. SNAP-MS analysis displayed in Cytoscape. Raw data files for SIRIUS analysis. Supplementary dataset 1. The CAZymes annotated result.

## Author Contributions

THe, XLi and KH designed the study and developed the workflow. THe designed and carried out the genomic analysis. XLi designed and performed the metabolomics analysis with contributions from AMR. RdCFV and JMvD designed and conducted antibacterial activity assays.KH supervised the project.

## Institutional Review Board Statement

Not applicable.

## Informed Consent Statement

Not applicable.

## Data Availability Statement

The sequencing data and genome assembly were deposited in the European Nucleotide Archive (ENA) at EMBL-EBI under accessions ERR14081000 and ERZ24985901, respectively. The mass spectrometry data were deposited on GNPS under the accession number MassIVE ID: MSV000095645.

## Acknowledgments

The authors are grateful for technical assistance by Pieter Tepper, assistance with data curation and ENA upload by Thomas Hackl, and the staff at the interfaculty mass spectrometry center of the University Medical Center RUG/UMCG.

## Abbreviations

ACP: acyl-carrier protein
AT: acyl-transferase
BGCs: biosynthetic gene clusters
BLAST: basic local alignment search tool
CAZymes: carbohydrate-active enzymes
CFUs: colony-forming units
CYA: czapek yeast autolysate agar
CZA: czapek’s agar
DH: dehydrogenase
DMSO: dimethyl sulfoxide
GenSAS: genome sequence annotation Server
GNPS: global natural products social molecular networking
GO: gene ontology
ER: enoyl reductase
ESI: electrospray ionization
EtOAc: ethyl acetate
FA: formic acid
HDAC: histone deacetylase
HPLC: high performance liquid chromatography
HR-LC-MS/MS: high-resolution liquid chromatography-coupled tandem mass spectrometry
ITS: internal transcribed spacer
KR: β-ketoreductase
KS: β-ketosynthase
LR-LC-MS: low-resolution liquid chromatography mass spectrometry
LSU: large ribosomal subunit
MBC: minimum bactericidal concentration
MEA: malt extract agar
MET: methyltransferase
MHB: mueller hinton broth
MIC: minimum inhibitory concentration
ML: maximum-likelihood
MRSA: methicillin-resistant *Staphylococcus aureus*
MSSA: methicillin-sensitive *Staphylococcus aureus*
NCBI: National Center for Biotechnology Information
NMR: nuclear magnetic resonance spectroscopy
NRP: non-ribosomal peptide
NRPKS: nonreducing polyketide synthase
ONT: oxford nanopore technology
PDA: potato dextrose agar
PDB: potato dextrose broth
PH: procaine hydrochloride
PK: polyketide
PT: product template
Ripp: ribosomally synthesized and post-translationally modified peptide
SB: sodium butyrate
SDA: sabouraud dextrose agar
SMs: secondary metabolites
TE: thioesterase
TICs: total ion chromatograms
*TUB2*: beta-tubulin 2.

