## Supplementary Figures and Tables for "The endophytic fungus *Cosmosporella* sp. VM-42 from *Vinca minor* is a source of bioactive compounds with potent activity against drug-resistant bacteria"

**Table S1**. List of species in *Nectriaceae* from NCBI used for phylogenetic analysis.

**Table S2**. Quality metrics for isolated genomic DNA of *Cosmosporella sp.* VM-42.

**Table S3.** Molecular networking-based identification of secondary metabolites in *Cosmosporella* sp. VM-42.

**Table S4**. Biosynthetic gene clusters of *Cosmosporella* sp. VM-42 predicted by antiSMASH version 7.1.0.

**Figure S1.** Contigs of *Cosmosporella* sp. VM-42 visualized by Bandage.

**Figure S2**. Antimicrobial activity of *Cosmosporella* sp. VM-42.

**Figure S3.** Epigenetic manipulation of *Cosmosporella* sp. VM-42 with sodium butyrate (SB; A) and procaine hydrochloride (PH; B).

**Figure S4.** ^1^H NMR spectrum of compound **27** in DMSO-d6 (600 MHz).

**Figure S5.** ^13^C NMR spectrum of compound **27** in DMSO-d6 (150 MHz).

**Figure S6.** ^1^H-^1^H COSY spectrum of compound **27** in DMSO-d6.

**Figure S7.** HSQC spectrum of compound **27** in DMSO-d6.

**Figure S8.** HMBC spectrum of compound **27** in DMSO-d6.

**Figure S9**. Functional genome annotation of *Cosmosporella* sp. VM-42

**Figure S10.** Compounds produced by BGCs that were returned as the most similar entries in MiBIG for certain BGCs detected in the *Cosmosporella* sp. VM-42 genome based on antiSMASH analysis.

Figure S11. BGCs in the *Cosmosporella* sp. VM-42 genome compared to homologous BGCs and their proposed products.

**Note S1.** Detailed annotations of putatively identified other compounds in the molecular network.

**Note S2.** Other biosynthetic potential of *Cosmosporella* sp. VM-42.

Table S1. List of species in *Nectriaceae* from NCBI used for phylogenetic analysis.

| Species | Accession number | | |
| --- | --- | --- | --- |
|  | ITS | LSU | *tub2* |
| *Cosmospora vilior* G.J.S. 90-217 | JF832596 | JF832681 | JF832840 |
| *Cosmospora scruposae* G.J.S. 86-320 | KJ676163 | KJ676200 | KJ676279 |
| *Cosmospora stilbohypoxyli* A.R. 4783 | KJ676144 | KJ676181 | KJ676257 |
| *Cosmospora ustulinae* MAFF 241532 | KJ676175 | KJ676212 | KJ676296 |
| *Cosmospora clavi* CBS 123941 | KJ676149 | KJ676186 | KJ676262 |
| *Cosmospora viliuscula* G.J.S 10-114 | KJ676155 | KJ676192 | KJ676270 |
| *Cosmospora khandalensis* A.R. 4799 | KJ676146 | KJ676183 | KJ676259 |
| ***Cosmospora arxii* CBS 748.69** | **KM231819** | **KM231694** | **KM232089** |
| *Cosmospora viliuscula* strain G.J.S. 83-197 | KC291732 | KC291777 | KC291907 |
| *Cosmospora viridescens* CBS 102433 | KJ676148 | KJ676185 | KJ676261 |
| ***Cosmospora cymosa* CBS 762.69** | **HQ897828** | **KM231693** | **KM232087** |
| ***Cosmospora coccinea* CBS 341.70** | **MH859703** | **KM231692** | **KM232086** |
| *Cosmospora fomiticola* G.J.S. 83-194 | KJ676158 | KJ676195 | KJ676274 |
| *Cosmosporella olivacea* KUMCC 17-0321 | MH087212 | MH087214 | MH087216 |
| *Cosmosporella olivacea* KUMCC 18-0016 | MH087213 | MH087215 | MH087217 |
| *‘Nectria’ flavoviridis* IMI 338173 | KC291747 | KC291785 | KC291902 |
| *Cosmosporella obscura* MAFF 241484 | KC291719 | KC291788 | KC291903 |
| *Pseudocosmospora rogersonii* G.J.S. 10-296 | KC291727 | KC291774 | KC291917 |
| *Pseudocosmospora eutypellae* G.J.S. 10-248 | KC291722 | KC291772 | KC291911 |
| *Dialonectria episphaeria* CBS 125494 | MH863609 | KM231697 | KM232092 |
| *Dialonectria ullevolea* CBS 125493 | KM231821 | KM231696 | KM232091 |
| ***Fusicolla aquaeductuum* CBS 837.85** | **KM231823** | **KM231699** | **KM232094** |
| *Fusicolla matuoi* CBS 581.78 | KM231822 | KM231698 | KM232093 |

The type strains are indicated in bold. Abbreviations: **A.R.**: Amy Y. Rossman personal collection; **ATCC**: American Type Culture Collection, Manassas, Virginia, U.S.A.; **CBS**: Centraalbureau voor Schimmelcultures, Utrecht, The Netherlands; **G.J.S.**: Gary J. Samuels personal collection; **IMI**: CABI Bioservices, Egham, Surrey, UK; **MAFF**: The Ministry of Agriculture, Forestry and Fisheries, Tsukuba, Japan; **KUMCC**: Kunming Institute of Botany Culture Collection, China.

**Table S2**. Quality metrics for isolated genomic DNA of *Cosmosporella sp.* VM-42.

| Qubit conc. (ng/µl) | 474.5 |
| --- | --- |
| NanoDrop conc. (ng/µl) | 337.0 |
| Output (µg) | 5.7 |
| A260/280 ratio | 1.89 |
| A260/230 ratio | 2.11 |
| NanoDrop/Qubit conc. ratio | 1.4 |
| Qubit conc. after Circulomics (ng/µl) | 50.6 |
| Input for library preparation (µg) | 1 |
| Output after DNA repair and end-prep (µg) | 0.855 |
| Output after adapter ligation and clean-up (µg) | 0.653 |

**Table S3.** Molecular networking-based identification of secondary metabolites in *Cosmosporella* sp. VM-42.

| **Compound** | **Metabolite name** | **rt [min]** | **Experimental mass [*m*/*z*]** | **Adduct** | **Molecular formula** | **Molecular weight** | **Exact mass [*m*/*z*]** | **MZ ErrorPPM [ppm]** | **MS^2^ fragment ions [*m*/*z*]** | **Samples** | |
| --- | --- | --- | --- | --- | --- | --- | --- | --- | --- | --- | --- |
|  |  |  |  |  |  |  |  |  |  | **SB 10 mM** | **PH 10 mM** |
| **1** | W493B/acuminatum B | 20.76 (SB), 19.21 (PH) | 888.5451 | [M+H]^+^ | C_45_H_73_N_7_O_11_ | 888.12 | 888.5464, 888.5488 | 1.46 (SB), 4.16 (PH) | 781.4880, 739.4390, 729.4471, 600.4036, 594.3884, 576.3771, 484.3408, 477.2348, 466.3292, 448.3183, 406.2004, 405.2144, 395.2917, 377.2810, 324.2541, 306.2434, 295.1660, 271.1411, 200.1037, 143.0820, 101.0712 | + | + |
| **2** | W493A/acuminatum C | 20.50 (SB), 18.83 (PH) | 874.5295 | [M+H]^+^ | C_44_H_71_N_7_O_11_ | 874.09 | 874.5309, 874.5333 | 1.60 (SB), 4.34 (PH) | 695.4323, 693.4553, 612.4006, 600.4350, 594.3880, 586.3868, 576.3779, 484.3395, 466.3286, 448.3181, 395.2915, 377.2809, 324.2541, 306.2435, 271.1410, 200.1037, 143.0820, 101.0712 | + | + |
| **3** | 3-(22-(dodecan-2-yl)-6-(4-hydroxybenzyl)-18-(1-hydroxyethyl)-3,12,15-trimethyl-2,5,8,11,14,17,20-heptaoxo-1-oxa-4,7,10,13,16,19-hexaazacyclodocosan-9-yl)propanamide | 19.91 (SB), 17.66 (PH) | 846.4982 | [M+H]^+^ | C_42_H_67_N_7_O_11_ | 846.04 | 846.5002 (SB), 846.5015 (PH) | 2.36 (SB), 3.89 (PH) | 594.3848, 584.3751, 576.3738, 484.3436, 466.3291, 448.3176, 435.1892, 434.2034, 395.2917, 377.2811, 363.1679, 324.2541, 306.2436, 271.1411, 253.1192, 200.1038, 143.0820, 101.0713 | + | + |
| **4** | W493D | 20.39 (SB), 18.62 (PH) | 886.5295 | [M+H]^+^ | C_45_H_71_N_7_O_11_ | 886.10 | 886.5298 (SB), 886.5331 (PH) | 0.33 (SB), 4.06 (PH) | 574.3623, 482.3234, 477.2351, 446.3024, 434.2053, 393.2756, 375.2660, 322.2386, 304.2282, 295.1664, 271.1412, 200.1033, 143.0819, 132.1025, 101.0713 | + | + |
| **5** | acuminatum F | 20.61 (SB), 18.68 (PH) | 904.5401 | [M+H]^+^ | C_45_H_73_N_7_O_12_ | 904.12 | 904.5415 (SB), 904.5443 (PH) | 1.65 (SB), 4.75 (PH) | 594.3892, 576.3752, 493.2313, 484.3416, 466.3302, 448.3177, 395.2916, 377.2811, 324.2542, 311.1610, 306.2435, 271.1411, 200.1038, 143.0820, 101.0713 | + | + |
| **6** | W493C | 19.96 (SB), 17.92 (PH) | 860.5138 | [M+H]^+^ | C_43_H_69_N_7_O_11_ | 860.06 | 860.5156, 860.5172 | 2.09 (SB), 3.95 (PH) | 586.4156, 572.3724, 548.3433, 438.2981, 420.2863, 405.2144, 367.2600, 349.2497, 296.2228, 295.1664, 278.2125, 271.1409, 200.1038, 143.0820, 101.0712 | + | + |
| **7** | 3-((3S,6R,9S,12R,15S,18R,22S)-6-(3,4-dihydroxybenzyl)-22-((R)-dodecan-2-yl)-18-((R)-1-hydroxyethyl)-3-isopropyl-12,15-dimethyl-2,5,8,11,14,17,20-heptaoxo-1-oxa-4,7,10,13,16,19-hexaazacyclodocosan-9-yl)propanamide | 20.21 (SB), 18.56 (PH) | 890.5244 | [M+H]^+^ | C_44_H_71_N_7_O_12_ | 890.09 | 890.5258 (SB), 890.5278 (PH) | 1.57 (SB), 3.81 (PH) | 594.3896, 576.3790, 484.3373, 479.2173, 466.3303, 448.3199, 407.1955, 395.2923, 377.2819, 324.2549, 306.2440, 297.1458, 271.1414, 200.1040, 152.0712, 143.0822, 118.0869, 101.0714 | + | + |
| **8** | 3-((3S,6R,9S,12R,15S,18R,22S)-22-((R,E)-dodec-5-en-2-yl)-6-(4-hydroxybenzyl)-18-((R)-1-hydroxyethyl)-3-isopropyl-12,15-dimethyl-2,5,8,11,14,17,20-heptaoxo-1-oxa-4,7,10,13,16,19-hexaazacyclodocosan-9-yl)propanamide | 20.07 (SB), 17.97 (PH) | 872.5138 | [M+H]^+^ | C_44_H_69_N_7_O_11_ | 872.07 | 872.5154 (SB), 872.5179 (PH) | 1.83 (SB), 4.69 (PH) | 574.3671, 482.3228, 464.3130, 446.3032, 434.2036, 393.2762, 375.2654, 346.1409, 322.2386, 304.2280, 281.1505, 271.1411, 200.1038, 143.0820, 118.0866, 101.0713 | + | + |
| **9** | 3-(22-(heptan-2-yl)-6-(4-hydroxybenzyl)-18-(1-hydroxyethyl)-3-isopropyl-12,15-dimethyl-2,5,8,11,14,17,20-heptaoxo-1-oxa-4,7,10,13,16,19-hexaazacyclodocosan-9-yl)propanamide | 17.52 (SB), 14.70 (PH) | 804.4512 | [M+H]^+^ | C_39_H_61_N_7_O_11_ | 803.96 | 804.4534 (SB), 804.4547 (PH) | 2.73 (SB), 4.35 (PH) | 516.3085, 506.2964, 463.2214, 414.2623, 396.2505, 391.2004, 378.2410, 343.2241, 325.2138, 307.2030, 281.1513, 271.1414, 254.1763, 236.1658, 143.0822, 118.0868, 101.0715 | + | + |
| **10** | 3-(22-(decan-2-yl)-6-(4-hydroxybenzyl)-18-(1-hydroxyethyl)-3-isopropyl-12,15-dimethyl-2,5,8,11,14,17,20-heptaoxo-1-oxa-4,7,10,13,16,19-hexaazacyclodocosan-9-yl)propanamide | 19.61 (SB), 17.08 (PH) | 846.4982 | [M+H]^+^ | C_42_H_67_N_7_O_11_ | 846.04 | 846.4997 (SB), 846.5019 (PH) | 1.77 (SB), 4.37 (PH) | 666.3894, 584.3693, 572.3988, 548.3491, 463.2206, 445.2096, 438.2972, 420.2876, 391.1996, 367.2603, 349.2497, 296.2229, 278.2125, 271.1412, 200.1039, 143.0820, 118.0867, 101.0713 | + | + |
| **11** | Cephalochromin/ ustilaginoidin F | 19.63 (SB), 16.18 (PH) | 519.1296 | [M+H]^+^ | C_28_H_22_O_10_ | 518.47 | 519.1304 (SB), 519.1313 (PH) | 1.54 (SB), 3.27 (PH) | 477.0792, 260.0689 | + | + |
| **12** | Ustilaginoidin E / Ustilaginoidin E1/ chaetochromin C | 20.19 (SB), 17.13 (PH) | 533.1453 | [M+H]^+^ | C_29_H_24_O_10_ | 532.50 | 533.1459 (SB), 533.1467 (PH) | 1.12 (SB), 2.62 (PH) | 515.1358, 477.0833, 274.0848, 260.0693 | + | + |
| **13** | Chaetochromin B/ isochaetochromin B_2_/ ustilaginoidin D/ isochaetochromin A1 | 20.67 (SB), 18.04 (PH) | 547.1609 | [M+H]^+^ | C_30_H_26_O_10_ | 546.53 | 547.1614 (SB), 547.1627 (PH) | 0.91 (SB), 3.28 (PH) | 529.1538, 491.1012, 274.0847, 67.0548, 55.0547 | + | + |
| **14** | Ustilaginoidin G/ Dihydroisoustilaginoidin A | 19.77 (SB) | 517.1140 | [M+H]^+^ | C_28_H_20_O_10_ | 516.46 | 517.1147 (SB), 517.1152 (PH) | 1.35 (SB) | 499.1058, 475.1084, 457.0545, 260.0690, 258.0533 | + | - |
| **15** | Ustilaginoidin A/isoustilaginoidin A | 15.15 (SB), 11.56 (PH) | 515.0983 | [M+H]^+^ | C_28_H_18_O_10_ | 514.44 | 515.0994 (SB), 515.1003 (PH) | 2.13 (SB), -3.88 (PH) | 515.0992, 487.1037, 473.0520 | + | + |
| **16** | Ustilaginoidin P/ Ustilaginoidin L | 16.23 (SB), 12.33 (PH) | 529.1140 | [M+H]^+^ | C_29_H_20_O_10_ | 528.47 | 529.1147 (SB), 529.1158 (PH) | 1.32 (SB), 3.40 (PH) | 501.1192, 487.0672, 473.0516, 431.0024 | + | + |
| **17** | R-de-O-methyllasiodiplodin | 15.10 (SB), 9.69 (PH) | 279.1602 | [M+H]^+^ | C_16_H_22_O_4_ | 278.35 | 279.1601 (SB), 279.1601 (PH) | -0.30 (SB), -0.30 (PH) | 261.1501, 243.1391, 233.1548, 221.1181, 219.1385, 205.0869, 193.0866, 179.0711, 177.0555, 165.0555, 153.0553, 139.0395, 137.0603, 125.0602, 85.0651, 69.0703 | + | + |
| **18** | Lasiodiplodin | 11.73 (SB), 10.53 (PH) | 293.1758 | [M+H]^+^ | C_17_H_24_O_4_ | 292.38 | 293.1757 (SB), 293.1757 (PH) | -0.45 (SB), -0.45 (PH) | 275.1653, 257.1546, 247.1706, 233.1546, 217.1228, 203.1073, 191.0711, 175.0762, 163.0761, 153.0554, 151.0758, 137.0602, 95.0858, 83.0858, 69.0702 | + | + |
| **19** | Dihydroresorcylide | 12.32 (SB), 9.42 (PH) | 293.1394 | [M+H]^+^ | C_16_H_20_O_5_ | 292.33 | 293.1393 (SB), 293.1393 (PH) | -0.5 (SB), -0.50 (PH) | 275.1285, 257.1185, 247.1338, 239.1069, 235.0965, 233.1183, 229.1233, 219.1381, 215.1079, 207.1029, 205.0863, 195.0663, 193.0506, 191.0712, 179.0710, 177.0554, 167.0714, 153.0552, 139.0394, 127.1123, 125.0969, 123.1172, 111.0809, 109.1016, 101.0601, 99.0809, 97.0651, 85.0650, 83.0857, 69.0702, 55.0548 | + | + |
| **20** | (E)-9-etheno-de-O-methyllasiodiplodin | 11.93 (SB), 9.07 (PH) | 291.1601 | [M+H]^+^ | C_17_H_22_O_4_ | 290.36 | 291.1600 (SB), 291.1602 (PH) | -0.63 (SB), 0.06 (PH) | 273.1496, 263.1659, 255.1397, 245.1545, 237.1116, 231.1393, 219.1024, 205.0868, 193.0866, 191.0707, 177.0917, 167.1073, 165.0557, 163.0760, 149.0602, 137.0603, 123,1169, 101.0596, 97.0651, 85.0652, 83.0857, 73.0648, 71.0494, 69.0701, 57.0338, 55.0547 | + | + |
| **21** | 7,12-dihydroxy-14-methoxy-3-methyl-3,4,5,6,7,8,9,10-octahydro-1H-benzo[c][1]oxacyclododecin-1-one | 11.34 (SB), 9.72 (PH) | 309.1707 | [M+H]^+^ | C_17_H_24_O_5_ | 308.37 | 309.1707 (SB), 309.1707 (PH) | 0 (SB), 0 (PH) | 291.1610, 273.1492, 263.1289, 251.1278, 249.1120, 247.1703, 245.1562, 233.1181, 221.1177, 205.1224, 195.1017, 193.0863, 181.0861, 169.0863, 153.0555, 127.1121, 123.0809, 111.0445, 109.1019, 101.0600, 99.0446, 95.0860, 81.0701, 69.0701, 59.0497, 57.0342 | + | + |
| **22** | [4]-gingerol | 12.43 (PH) | 249.1496 | [M- H_2_O+ H]^+^ | C_15_H_22_O_4_ | 266.34 | 249.1496 (PH) | 0 (PH) | 249.1497, 231.1391, 207.1024, 205.1235, 193.0867, 179.0711, 177.0720, 169.0858, 163.0762, 151.0761, 137.0604, 125.0970, 111.0445, 109.1017, 97.0654, 95.0860, 87.0807, 85.0647, 73.0650, 69.0703, 55.0548, | - | + |
| **23** | [6]-gingerol | 14.6 (SB), 11.47 (PH) | 277.1809 | [M- H_2_O+ H]^+^ | C_17_H_26_O_4_ | 294.39 | 277.1804 (SB), 277.1806 (PH) | -1.87 (SB), -1.15 (PH) | 277.1811, 259.1700, 207.1022, 193.0865, 189.0919, 181.0869, 179.0713, 177.0919, 163.0759, 151.0760, 137.0603, 123.0806, 97.1016, 85.0651, 83.0858, 71.0495 | + | + |
| **24** | 4-(5-hydroxy-3-oxodecyl)-2-methoxyphenyl hydrogen sulfate | 13.86 (SB), 10.44 (PH) | 357.1377 | [M- H_2_O+ H]^+^ | C_17_H_26_O7S | 374.45 | 357.1379 (SB), 357.1381 (PH) | 0.47 (SB), 1.03 (PH) | 357.1388, 323.1503, 277.1812, 259.1705, 231.0332, 139.1123, 125.0970, 123.0807, 115.0762, 107.0858, 101.0601, 97.1016, 85.0652, 83.0859, 71.0495, 57.0705 | + | + |
| **25** | 2-(3-carboxypropanoylamino)-3-indol-3-ylpropanoic acid | 8.95 (SB), 6.12 (PH) | 305.1142 | [M+H]^+^ | C_15_H_16_N_2_O_5_ | 304.30 | 305.1141 (SB), 305.1144 (PH) | -0.32 (SB), 0.65 (PH) | 259.1088, 241.0981, 205.0981, 188.0715, 159.0924, 130.0655, 101.0238 | + | + |
| **26** | PyroGlu-Phe | 6.62 (SB), 5.26 (PH) | 277.1193 | [M+H]^+^ | C_14_H_16_N_2_O_4_ | 276.29 | 277.1189 (SB), 277.1193 (PH) | -1.44 (SB), 0 (PH) | 259.1061, 231.1137, 203.1188, 120.0813, 84.0446 | + | + |

Table S3. Biosynthetic gene clusters of *Cosmosporella* sp. VM-42 predicted by antiSMASH version 7.1.0.

| **Cluster ID** | **Region** | **Contig** | **Type** | **From (nt)** | **To (nt)** | **Most similar known cluster** | **Type** | **Similarity** |
| --- | --- | --- | --- | --- | --- | --- | --- | --- |
| 1 | 5.1 | 7 | [isocyanide-nrp](https://docs.antismash.secondarymetabolites.org/glossary/#isocyanide-nrp) | 293,146 | 348,355 |  |  |  |
| 2 | 5.2 | 7 | [fungal-RiPP-like](https://docs.antismash.secondarymetabolites.org/glossary/#fungal-ripp-like) | 2,887,943 | 2,949,096 |  |  |  |
| 3 | 5.3 | 7 | [terpene](https://docs.antismash.secondarymetabolites.org/glossary/#terpene) | 3,465,651 | 3,486,692 |  |  |  |
| 4 | 6.1 | 19 | [T1PKS](https://docs.antismash.secondarymetabolites.org/glossary/#t1pks) | 164,837 | 212,594 |  |  |  |
| 5 | 6.2 | 19 | [NRPS-like](https://docs.antismash.secondarymetabolites.org/glossary/#nrps-like) | 1,852,875 | 1,896,637 |  |  |  |
| 6 | 6.3 | 19 | [terpene](https://docs.antismash.secondarymetabolites.org/glossary/#terpene) | 4,100,761 | 4,122,304 | squalestatin S1 | Terpene | 40% |
| 7 | 7.1 | 5 | [NRPS](https://docs.antismash.secondarymetabolites.org/glossary/#nrps) | 2,004,181 | 2,050,034 | metachelin C/metachelin A/metachelin A-CE/metachelin B/dimerumic acid 11-mannoside/dimerumic acid | Polyketide | 62% |
| 8 | 7.2 | 5 | [NRPS](https://docs.antismash.secondarymetabolites.org/glossary/#nrps),[T1PKS](https://docs.antismash.secondarymetabolites.org/glossary/#t1pks) | 2,973,260 | 3,028,878 | UNII-YC2Q1O94PT | Polyketide | 100% |
| 9 | 9.1 | 12 | [T1PKS](https://docs.antismash.secondarymetabolites.org/glossary/#t1pks) | 1,468,521 | 1,510,510 | betaenone A/betaenone B/betaenone C | Polyketide | 25% |
| 10 | 9.2 | 12 | [T3PKS](https://docs.antismash.secondarymetabolites.org/glossary/#t3pks) | 1,682,115 | 1,723,580 |  |  |  |
| 11 | 9.3 | 12 | [T1PKS](https://docs.antismash.secondarymetabolites.org/glossary/#t1pks) | 1,756,417 | 1,804,460 |  |  |  |
| 12 | 9.4 | 12 | [terpene](https://docs.antismash.secondarymetabolites.org/glossary/#terpene) | 2,758,827 | 2,779,921 |  |  |  |
| 13 | 9.5 | 12 | [terpene](https://docs.antismash.secondarymetabolites.org/glossary/#terpene) | 2,850,875 | 2,872,756 |  |  |  |
| 14 | 9.6 | 12 | [T1PKS](https://docs.antismash.secondarymetabolites.org/glossary/#t1pks) | 2,972,247 | 3,019,737 |  |  |  |
| 15 | 9.7 | 12 | [NRPS-like](https://docs.antismash.secondarymetabolites.org/glossary/#nrps-like) | 3,207,757 | 3,250,873 |  |  |  |
| 16 | 9.8 | 12 | [T1PKS](https://docs.antismash.secondarymetabolites.org/glossary/#t1pks) | 3,369,626 | 3,417,416 |  |  |  |
| 17 | 9.9 | 12 | [NRPS](https://docs.antismash.secondarymetabolites.org/glossary/#nrps) | 3,581,698 | 3,625,457 |  |  |  |
| 18 | 11.1 | 4 | [T1PKS](https://docs.antismash.secondarymetabolites.org/glossary/#t1pks),[isocyanide](https://docs.antismash.secondarymetabolites.org/glossary/#isocyanide) | 59,991 | 137,684 |  |  |  |
| 19 | 11.2 | 4 | [NRPS-like](https://docs.antismash.secondarymetabolites.org/glossary/#nrps-like) | 159,209 | 202,523 |  |  |  |
| 20 | 11.3 | 4 | [T1PKS](https://docs.antismash.secondarymetabolites.org/glossary/#t1pks) | 321,144 | 390,568 | azasperpyranone A/azasperpyranone B/azasperpyranone C/azasperpyranone D/azasperpyranone E/azasperpyranone F/azasperpyranone G/azasperpyranone H | Polyketide | 18% |
| 21 | 11.4 | 4 | [NRPS](https://docs.antismash.secondarymetabolites.org/glossary/#nrps),[T1PKS](https://docs.antismash.secondarymetabolites.org/glossary/#t1pks) | 3,582,742 | 3,672,868 | W493 B/W493 A | NRP+Polyketide | 100% |
| 22 | 12.1 | 3 | [terpene](https://docs.antismash.secondarymetabolites.org/glossary/#terpene) | 3,339,684 | 3,360,829 |  |  |  |
| 23 | 12.2 | 3 | [NRPS](https://docs.antismash.secondarymetabolites.org/glossary/#nrps) | 5,447,121 | 5,493,428 |  |  |  |
| 24 | 13.1 | 15 | [NRPS](https://docs.antismash.secondarymetabolites.org/glossary/#nrps) | 149,157 | 195,581 | [gliovirin](https://mibig.secondarymetabolites.org/go/BGC0001609/1) | NRP | 25% |
| 25 | 13.2 | 15 | [T1PKS](https://docs.antismash.secondarymetabolites.org/glossary/#t1pks) | 211,421 | 259,475 | [nectriapyrone C/nectriapyrone D/nectriapyrone](https://mibig.secondarymetabolites.org/go/BGC0002155/1) | Polyketide | 100% |
| 26 | 13.3 | 15 | [T1PKS](https://docs.antismash.secondarymetabolites.org/glossary/#t1pks) | 2,682,418 | 2,727,740 | [ustilaginoidin N/ustilaginoidin O/ustilaginoidin M/ustilaginoidin A/ustilaginoidin F/ustilaginoidin E/ustilaginoidin D/ustilaginoidin G](https://mibig.secondarymetabolites.org/go/BGC0002177/1) | Polyketide | 46% |
| 27 | 14.1 | 9 | [NRPS-like](https://docs.antismash.secondarymetabolites.org/glossary/#nrps-like) | 1,129,132 | 1,173,429 |  |  |  |
| 28 | 14.2 | 9 | [T1PKS](https://docs.antismash.secondarymetabolites.org/glossary/#t1pks) | 3,020,490 | 3,067,156 | [HEx-pks23 polyketide](https://mibig.secondarymetabolites.org/go/BGC0002215/1) | Polyketide | 44% |
| 29 | 15.1 | 1 | [NRPS](https://docs.antismash.secondarymetabolites.org/glossary/#nrps),[terpene](https://docs.antismash.secondarymetabolites.org/glossary/#terpene) | 203,431 | 259,434 |  |  |  |
| 30 | 15.2 | 1 | [terpene](https://docs.antismash.secondarymetabolites.org/glossary/#terpene) | 1,346,435 | 1,367,724 |  |  |  |
| 31 | 15.3 | 1 | [betalactone](https://docs.antismash.secondarymetabolites.org/glossary/#betalactone) | 3,451,146 | 3,478,897 |  |  |  |
| 32 | 16.1 | 2 | [NRPS-like](https://docs.antismash.secondarymetabolites.org/glossary/#nrps-like) | 569,698 | 613,549 | [choline](https://mibig.secondarymetabolites.org/go/BGC0002276/1) | NRP | 100% |
| 33 | 16.2 | 2 | [NRP-metallophore](https://docs.antismash.secondarymetabolites.org/glossary/#nrp-metallophore),[NRPS](https://docs.antismash.secondarymetabolites.org/glossary/#nrps) | 2,681,682 | 2,743,085 |  |  |  |
| 34 | 16.3 | 2 | [NRPS](https://docs.antismash.secondarymetabolites.org/glossary/#nrps) | 2,820,370 | 2,871,996 |  |  |  |
| 35 | 16.4 | 2 | [terpene](https://docs.antismash.secondarymetabolites.org/glossary/#terpene) | 3,525,482 | 3,547,836 |  |  |  |

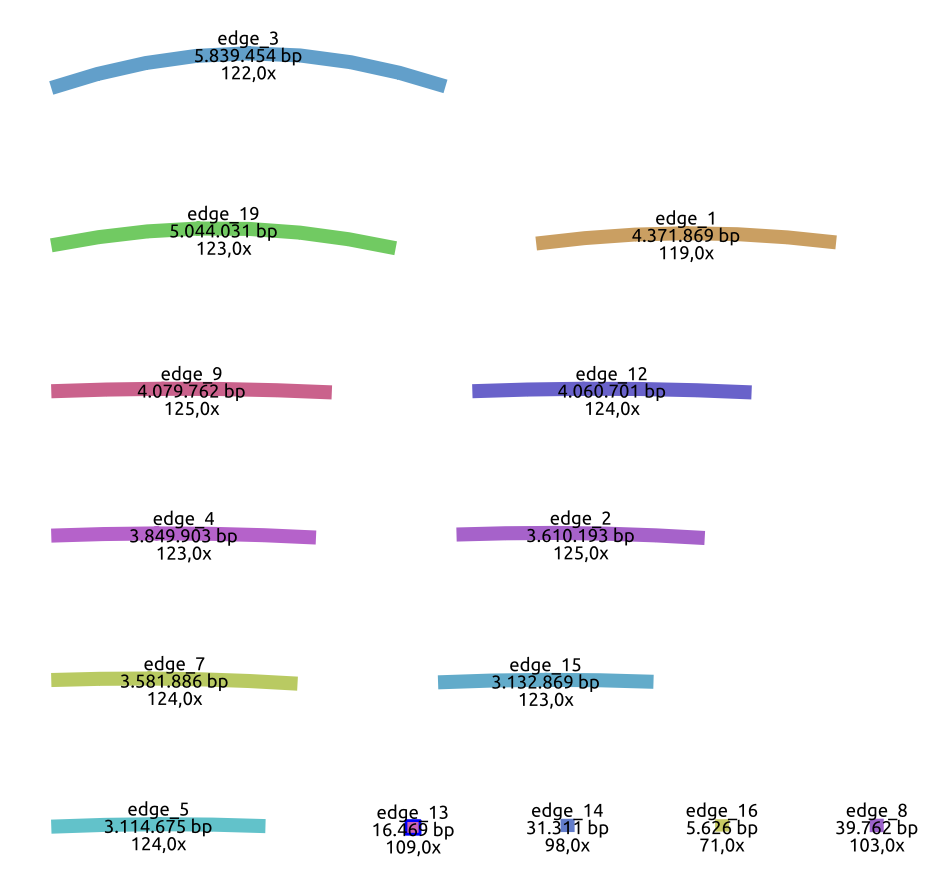

**Figure S1**. Contigs (was assembled by Flye with all the reads, including those from mitochondria) of *Cosmosporella* sp. VM-42 visualized by Bandage.

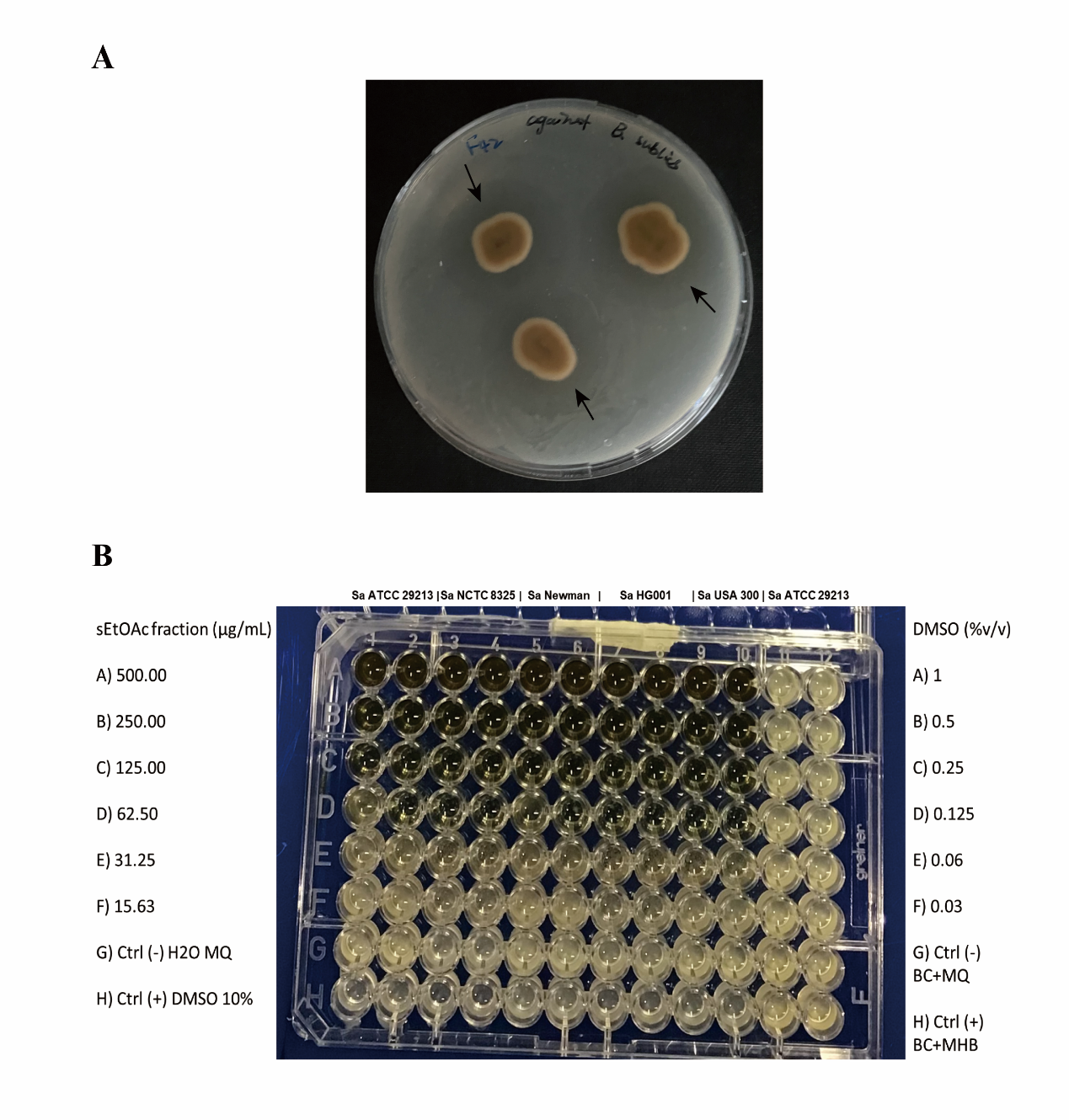

**Figure S2**. Antimicrobial activity of *Cosmosporella* sp. VM-42*.* (A) Agar diffusion test for antimicrobial activity of *Cosmosporella* sp. VM-42 against *B. subtilis.* The black arrows indicate growth inhibition zones around three fungal colonies; (B) 96-well plate showing the inhibitory activity of the sEtOAc extract of *Cosmosporella* sp. VM-42 against different MSSA and MRSA strains indicated on top of the image.

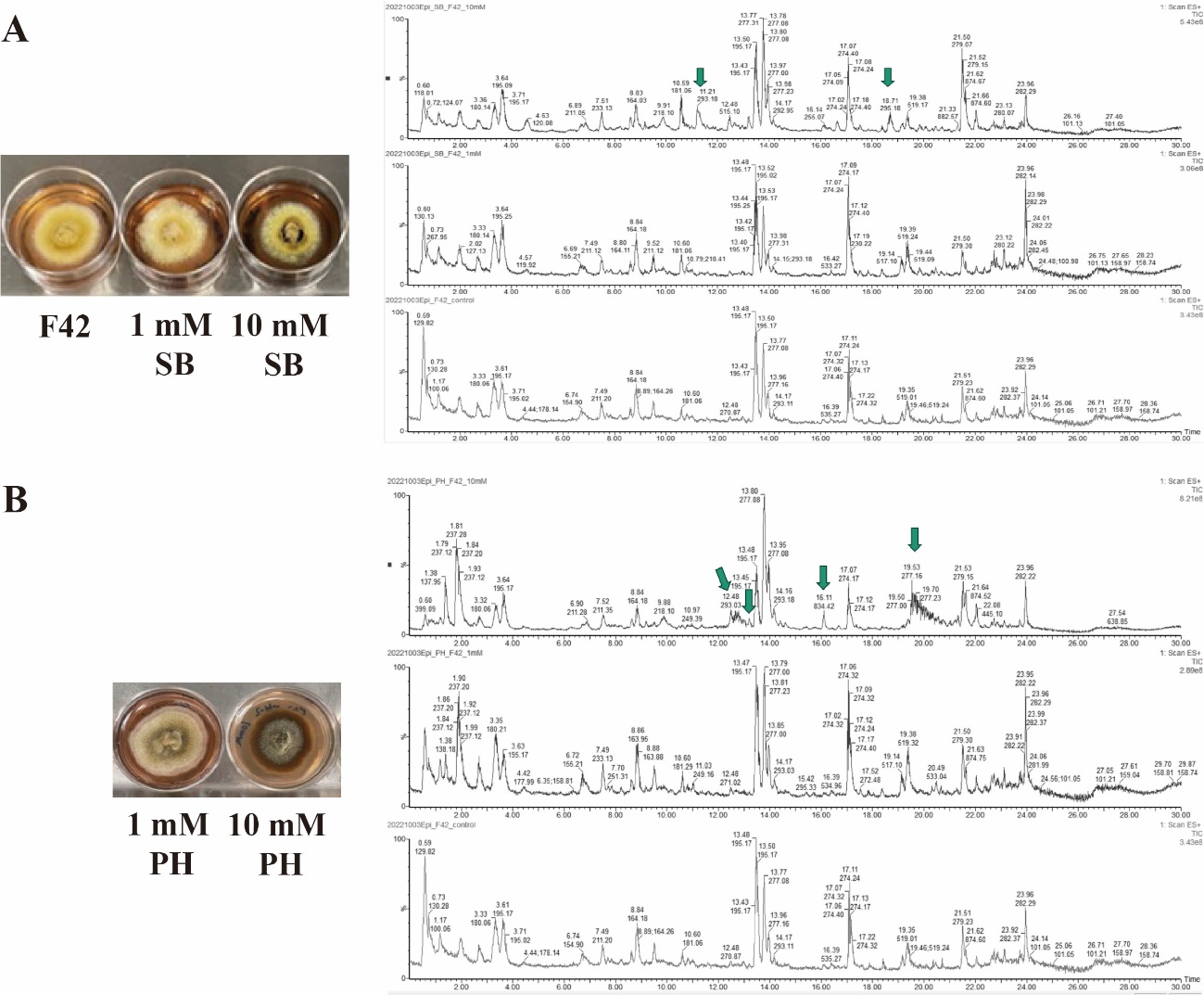

**Figure S3.** Epigenetic manipulation of *Cosmosporella* sp. VM-42 with sodium butyrate (SB; A) and procaine hydrochloride (PH; B). Colony morphology of the fungus treated with SB or PH and total ion chromatograms (TIC) recorded with low-resolution LC-MS of EtOAc crude extracts. The experiment was performed in triplicate, and outcomes were recorded after 14 days of cultivation at 25°C. Green arrows indicate newly observed or enhanced peaks following treatment with SB and PH.

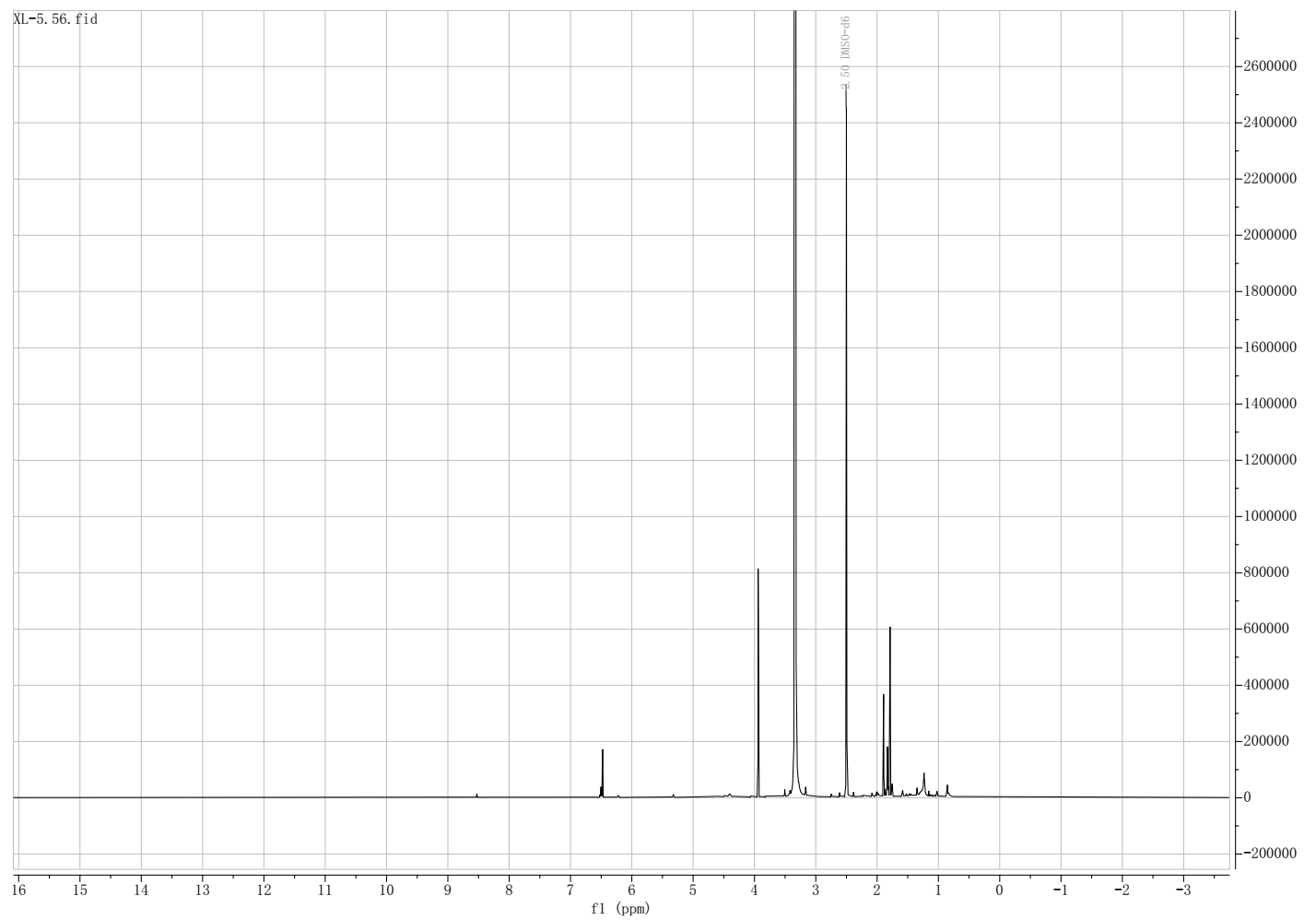

Figure S4. ^1^H NMR spectrum of compound 27 in DMSO-d6 (600 MHz).

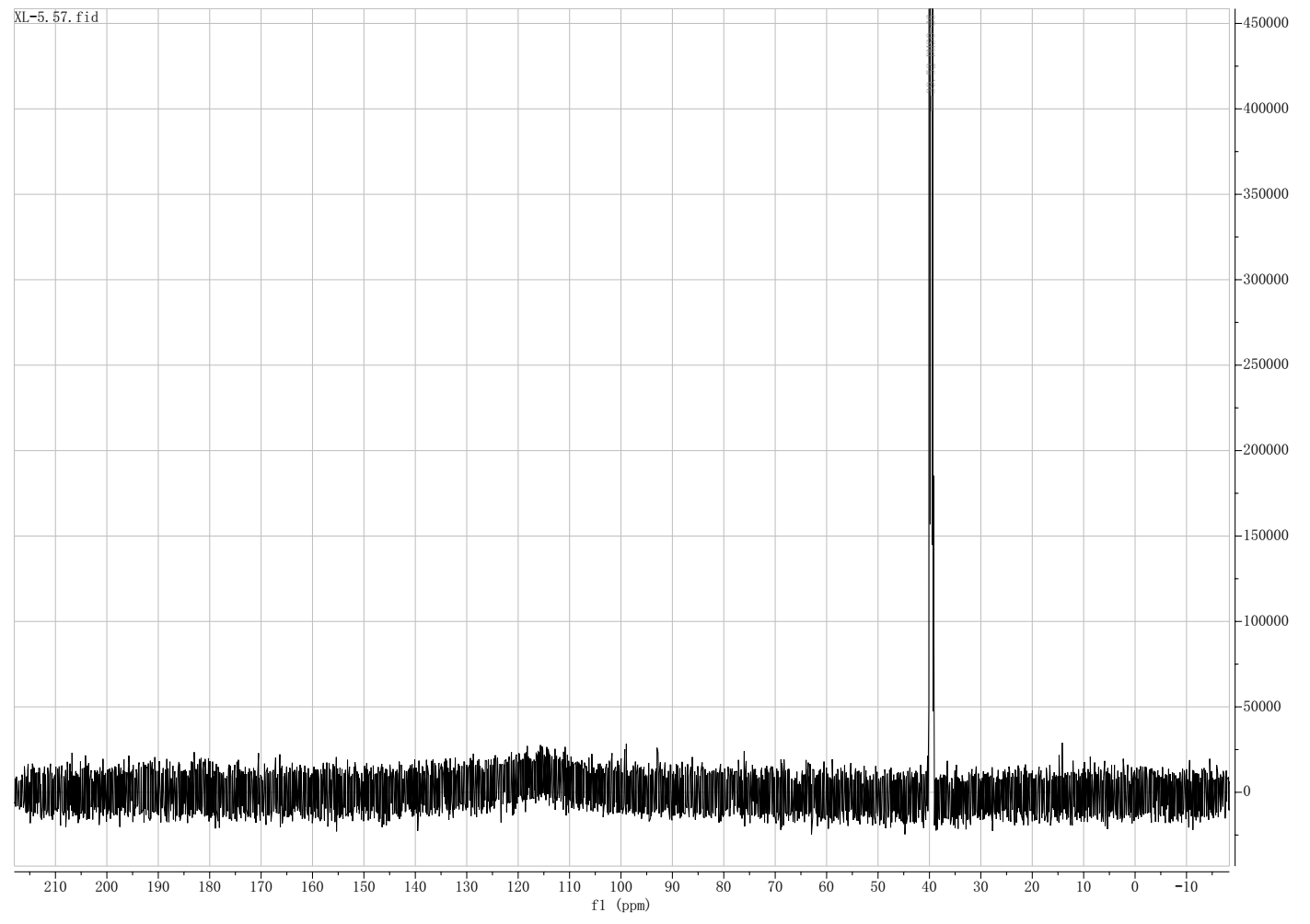

**Figure S5.** ^13^C NMR spectrum of compound **27** in DMSO-d6 (150 MHz).

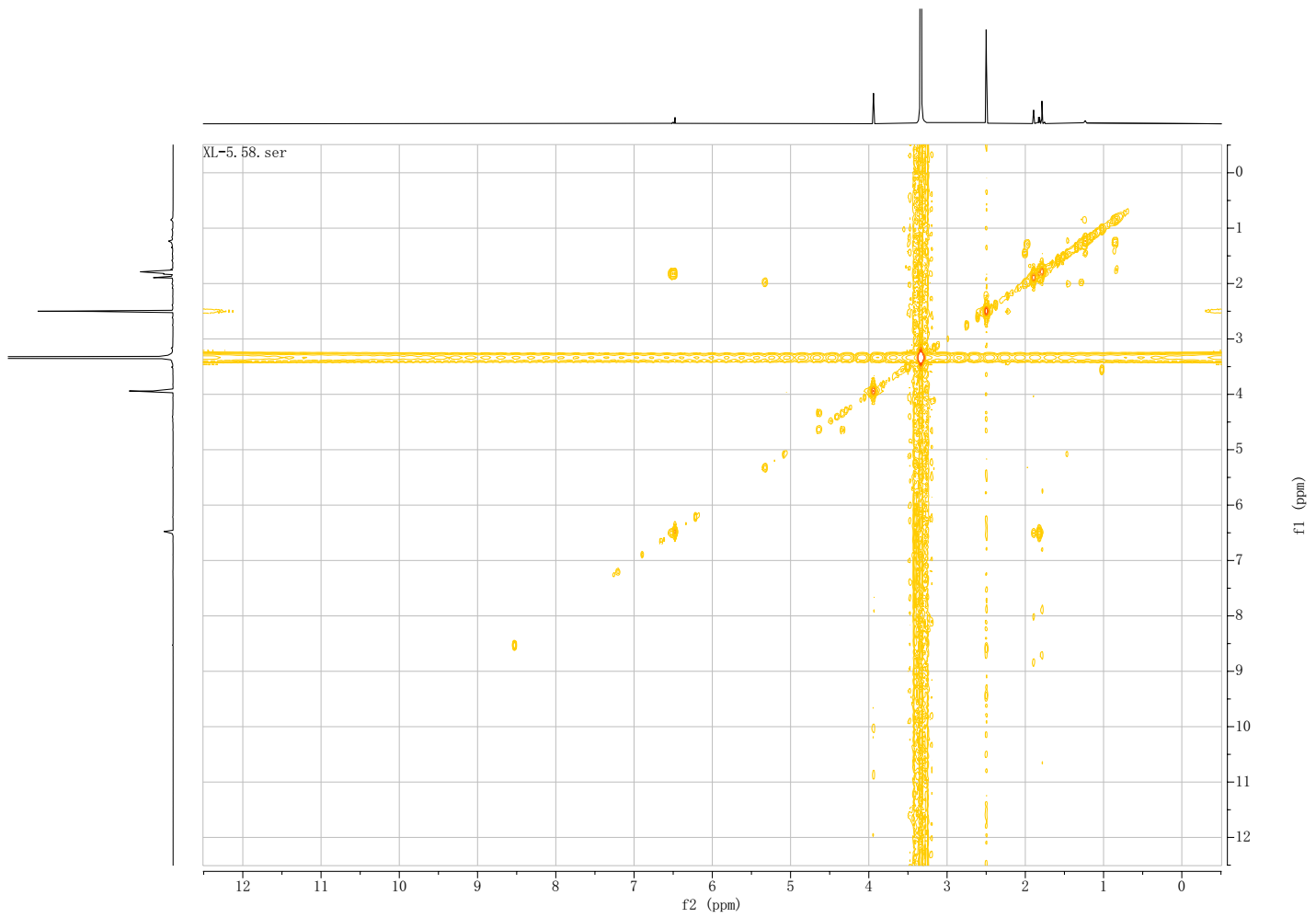

Figure S6. ^1^H-^1^H COSY spectrum of compound 27 in DMSO-d6.

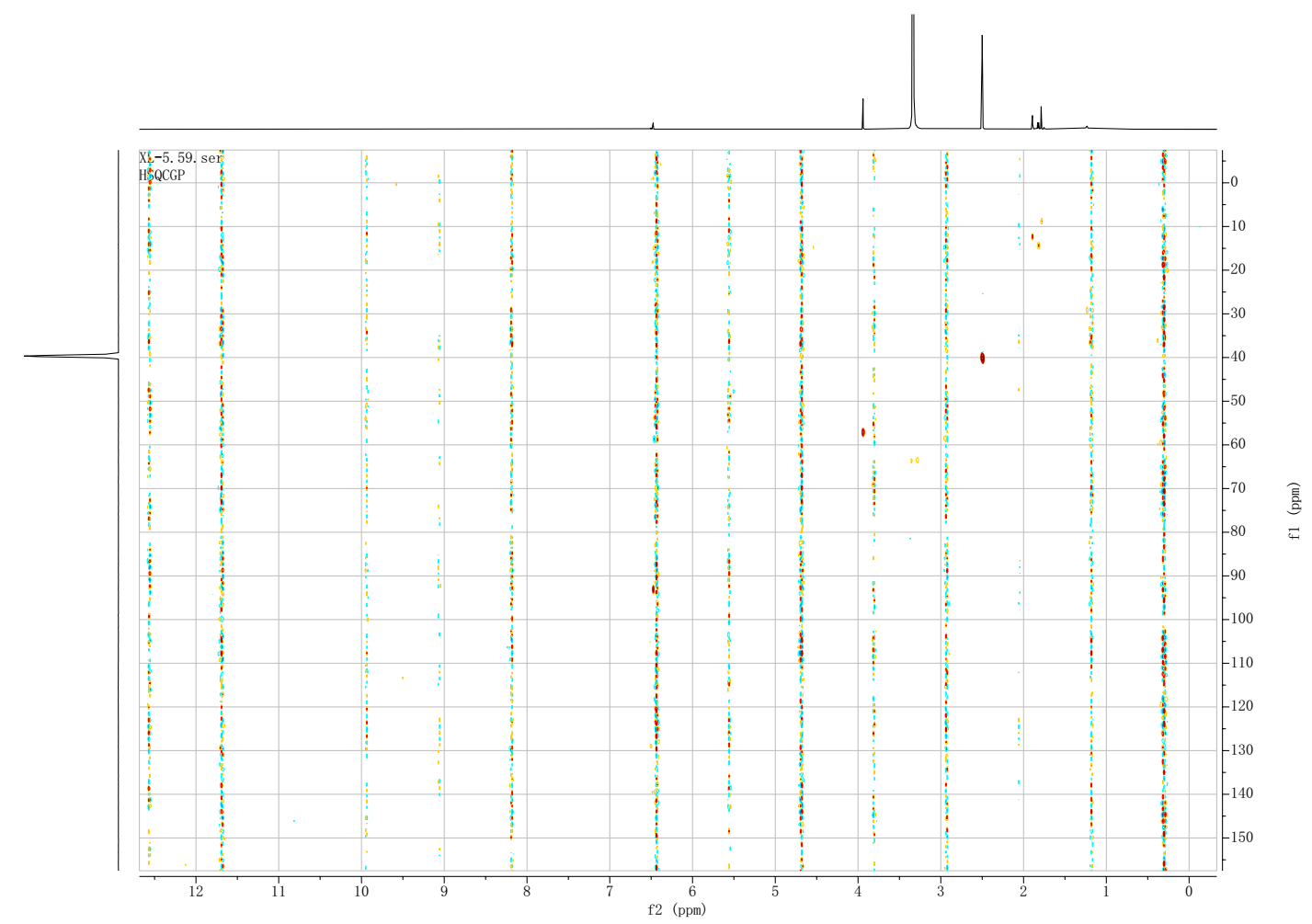

Figure S7. HSQC spectrum of compound 27 in DMSO-d6.

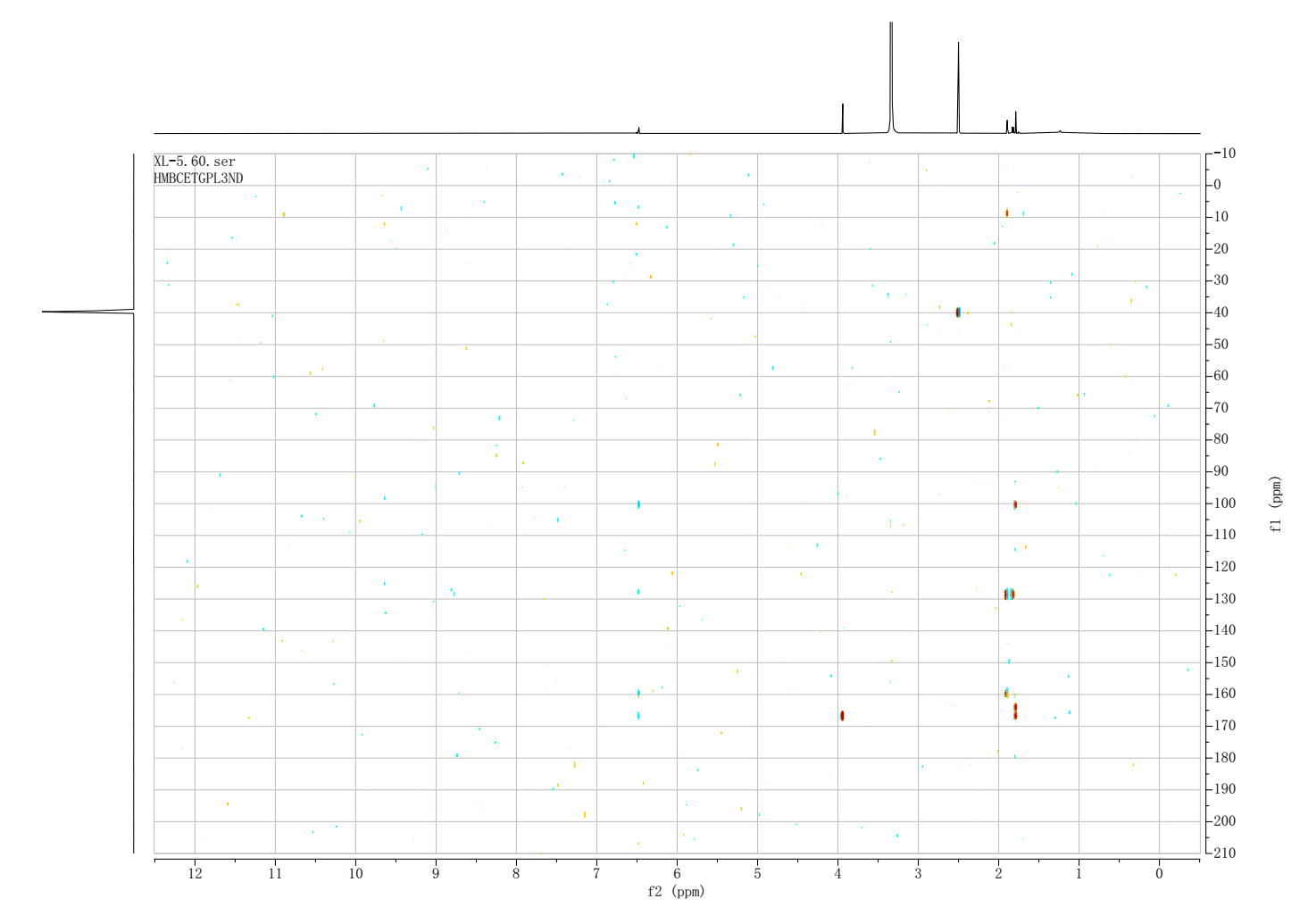

Figure S8. HMBC spectrum of compound 27 in DMSO-d6.

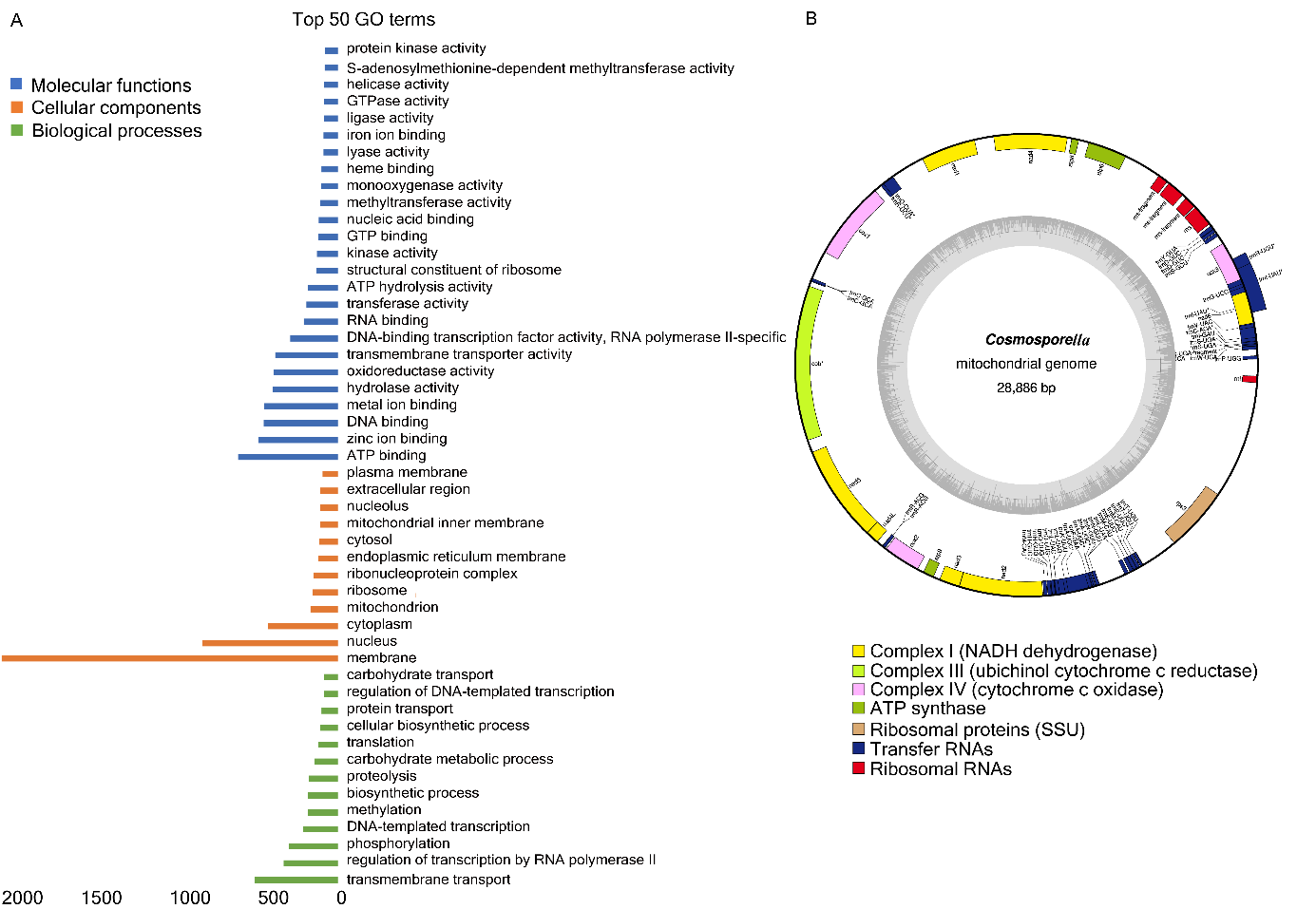

Figure S9. Functional genome annotation of *Cosmosporella* sp. VM-42*.*

(A) Gene Ontology (GO) analysis of the annotated *Cosmosporella* sp. VM-42 genome; top 50 terms were grouped into the three major GO terms “molecular functions” (50%), “biological processes” (26%), and “cellular components” (50%) (Figure S9A). In total, 415 genes were annotated as CAZymes in dbCAN, including 52% glycoside hydrolases (GHs), 22% glycosyl transferases (GTs), 15% auxiliary activities (AAs), 6% carbohydrate esterases (CEs), 3% polysaccharide lyases (PLs), and 1% carbohydrate-binding modules (CBMs) (Supplementary dataset 1). The most abundant (>12 counts) CAZyme types in *Cosmosporella* sp. VM-42 are GH16 (22), GH18 (19), GH3 (14), GH5 (13), GT2(16) AA7 (20), and CE3 (12).

(B) Complete structural and functional annotation of the assembled mitochondrial genome of *Cosmosporella* sp. VM-42.

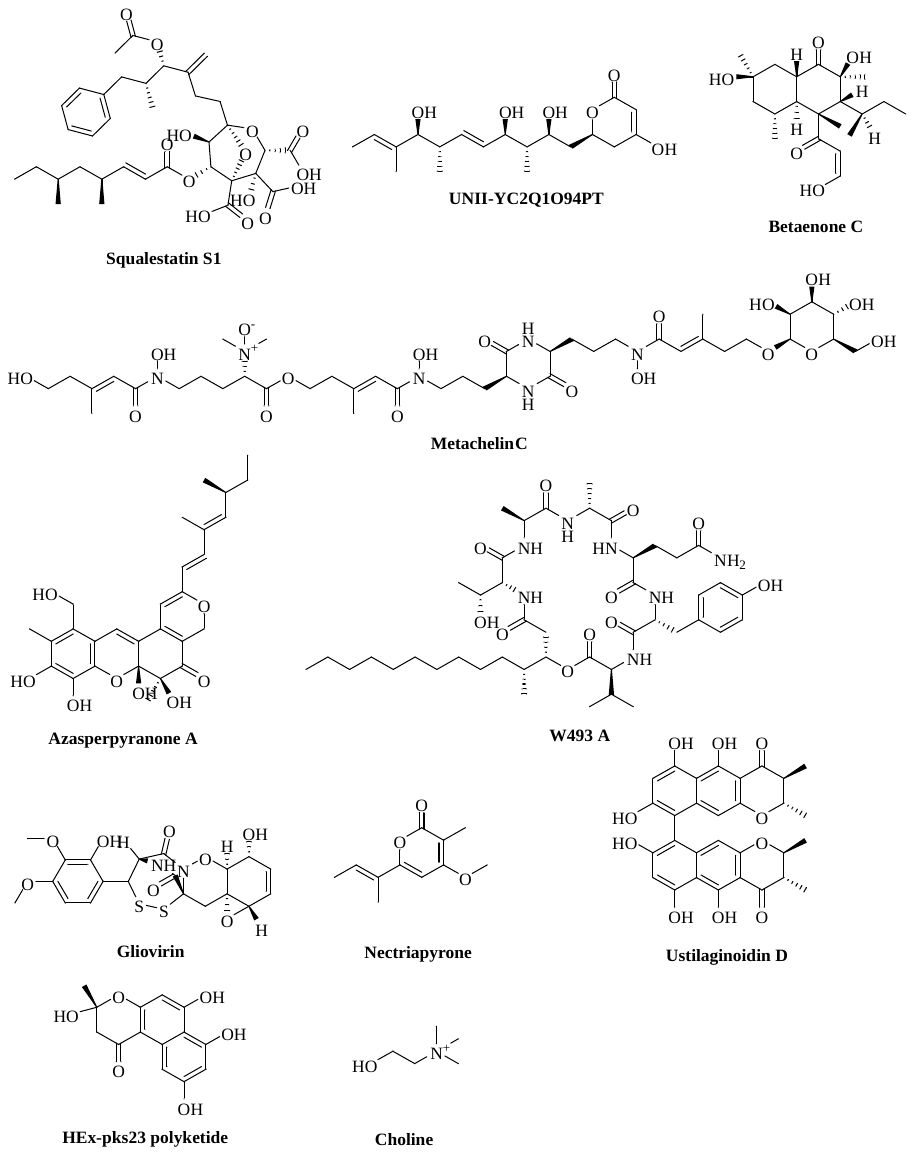
Figure S10. Compounds produced by BGCs that were returned as most similar entries in MiBIG for certain BGCs detected in the *Cosmosporella sp.* VM-42 genome based on antiSMASH analysis.

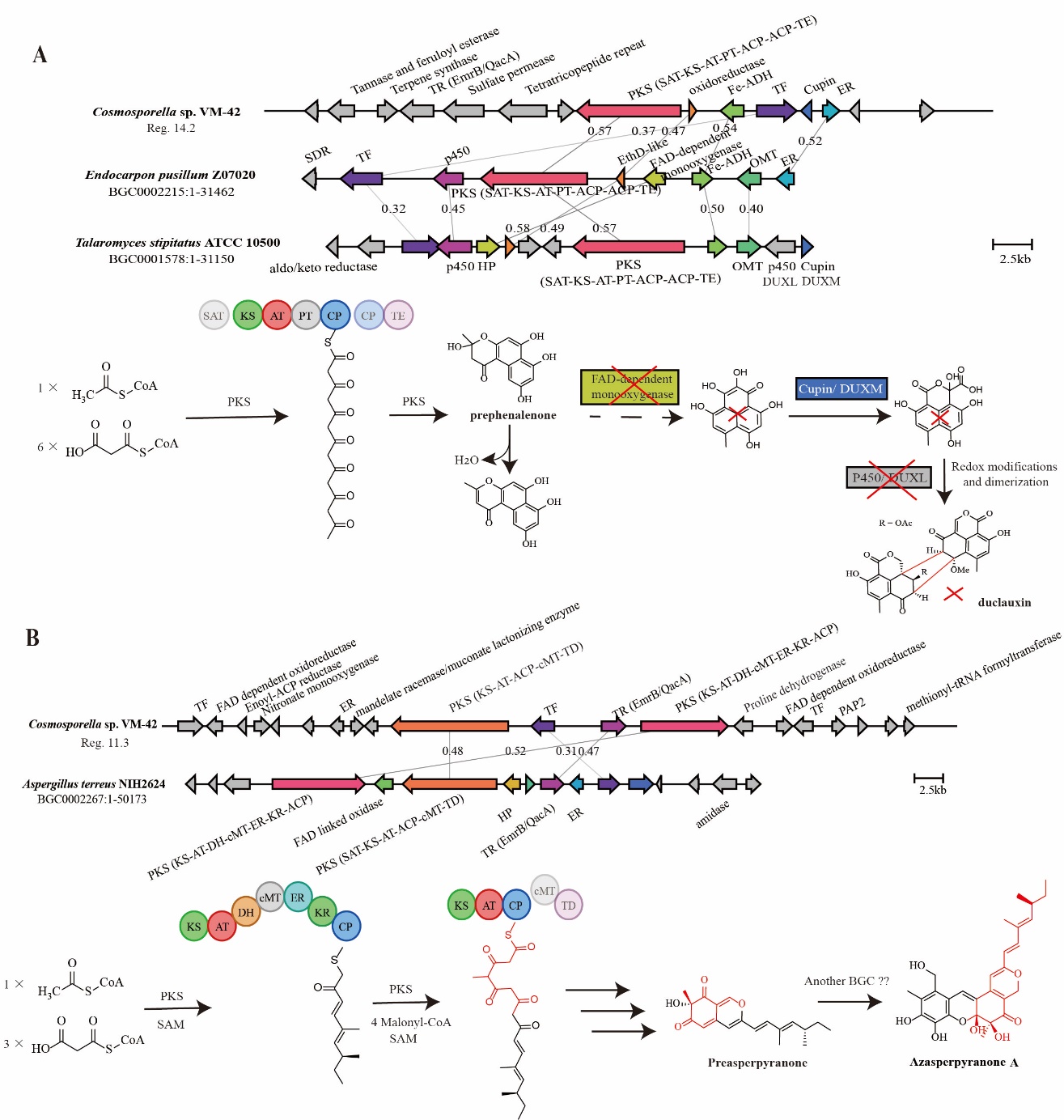

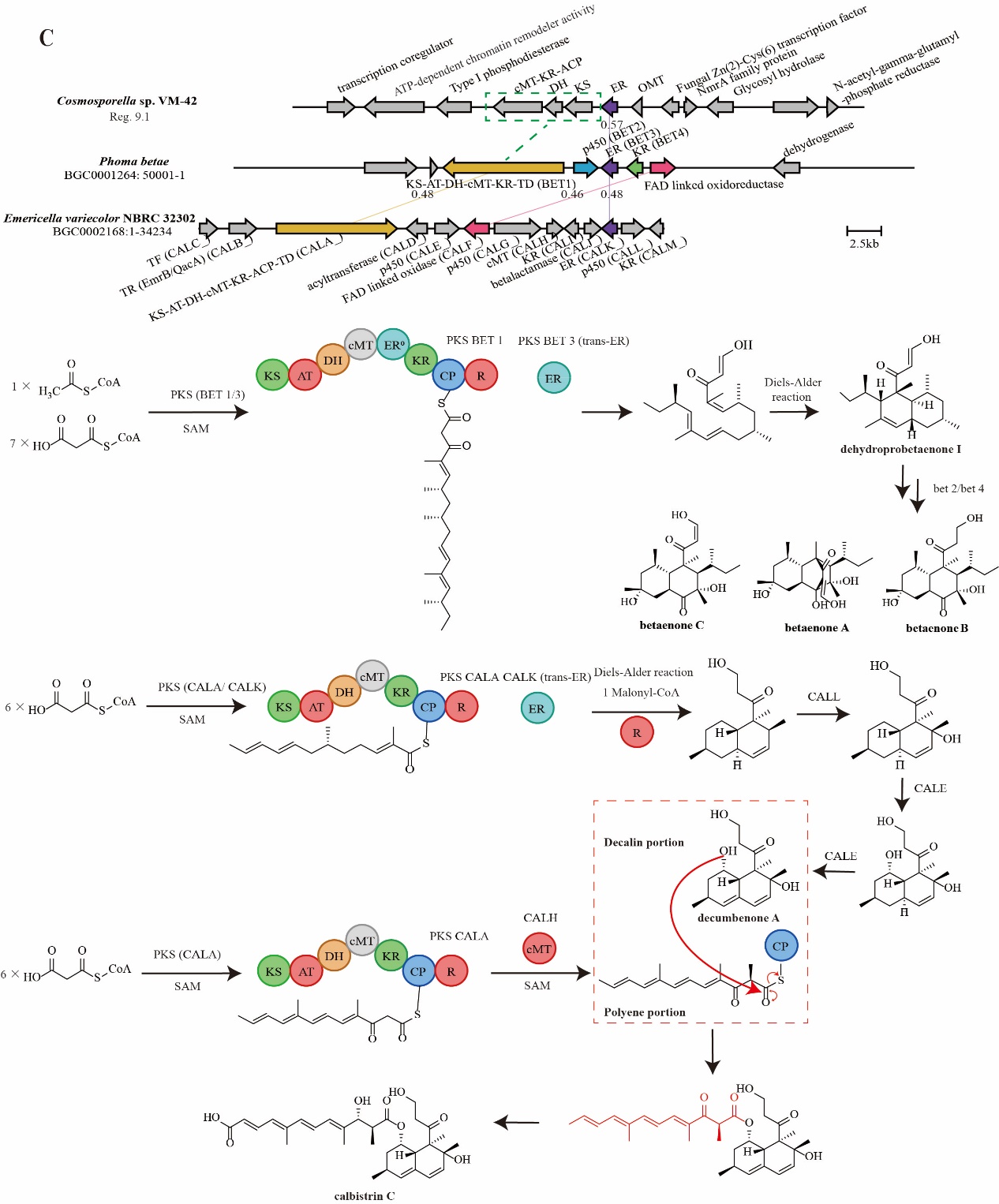

Figure S11. BGCs in the *Cosmosporella* sp. VM-42 genome compared to homologous BGCs and their proposed products. (A) Region 14.2 is homologous to BGCs from *Endocarpon pusilum* Z07020 and *Talaromyces stipitatus* ATCC 10500; (B) Region 11.3 is homologous to a BGC from *Aspergillus terreus* NIH2624; (C) Region 9.1 is homologous to BGCs from *Phoma betae* and *Emericella variecolor* NBRC 32302. Abbreviations: T, peptide acyl carrier domain; C, condensation domain; A, adenylation domain; E, epimerization; AMP, AMP binding; TR, transporter; TE, thioesterase; KS, ketosynthase; AT, acyltransferase; DH, dehydratase; cMT, C-methyltransferase; ER, ene-reductase; KR, ketoreductase; ACP, acyl carrier protein; Pep, peptidase; DUF, domain of unknown function; SAT, starter unit acyltransferase; PT, product template; Fe-ADH, Iron-containing alcohol dehydrogenase; HP, hypothetical protein; OMT, O-methyltransferase; EthD, EthD family protein; P450, cytochrome P450; FAD, flavin adenine dinucleotide; TF, transcription factor. Gene cluster comparison and visualization were performed by clinker on the CAGECAT web server (https://cagecat.bioinformatics.nl/). The small numbers on the lines connecting genes are amino acid sequence identities expressed in decimal fractions.

Note S1. Detailed annotations of putatively identified 26 compounds in the molecular network.

Cluster 2 has 3 nodes in total, which are identified as naphtho-γ-pyrones. The node with *m/z* 519.131 (**11**) was annotated as cephalochromin by GNPS, SNAP-MS and Sirius, which is a homodimer of hemiustilaginoidin F. Since the C-C bond of the dimer junction can freely rotate and due to the unknown stereochemistry of the substitutions, the three-dimensional configuration of the dimer cannot be determined based on MS2 results. Therefore, **11** can also be ustilaginoidin F, with the opposite configuration compared to cephalochromin. The node with *m/z* 533.146 (**12**) shows one -CH_2_ group more than **11**. Considering the results from SNAP-MS and Sirius analysis combined with unknown stereocenters on the compound’s structure, ustilaginoidin E, ustilaginoidin E1, chaetochromin C, or different configurations thereof, could be **12**. The node with *m/z* 547.162 (**13**) also has one -CH_2_ group more compared with **12**. Similarly, based on MS2 fragmentation information in Table 1, SNAP-MS and Sirius results, compound **13** was identified as chaetochromin B, isochaetochromin B_2_, ustilaginoidin D/ chaetochromin A, isochaetochromin A1, or the different configurations thereof. In addition to this cluster, a single node with *m/z* 517.115 (**14**), which presented fragments with *m/z* 499.1058, *m/z* 475.1084, *m/z* 457.0545, *m/z* 260.0690, and *m/z* 258.0533, is predicted to be ustilaginoidin G or dihydroisoustilaginoidin A by Sirius. Similarly, based on Sirius analysis, another node with *m/z* 515.0983 (**15**) in cluster 3 is predicted to be ustilaginoidin A or isoustilaginoidin A. According to its MS2 fragments, and its connected node with *m/z* 539.1147 (**16**) has one methyl group more than **15**, and is predicted to be ustilaginoidin L or ustilaginoidin P. These ustilaginoidins possess complex chirality, including not only the axial chirality, but also the stereogenic centers in the 2,3-dihydropyran-4-one ring when present (Lai et al., 2019; Sagita et al., 2021). Their absolute configuration can therefore only be determined by NMR, X-ray crystallography, or circular dichroism analyses of pure compounds (Lai et al., 2019).

Although cluster 4 has no GNPS annotations, SNAP-MS and Sirius analysis give us more information. The node with *m/z* 279.16 (**17**) is predicted to be an [M + H]^+^ adduct of R-de-O-methyllasiodiplodin, a type of macrolactone, and the node with *m/z* 261.14 represents the same structure with a different [M - H_2_O + H]^+^ adduct. The node with *m/z* 293.176 (**18**) has one -CH_2_ group more than **17**, and it is highly likely that the hydroxyl group will be converted to methoxy, which is predicted to be an [M + H]^+^ adduct of lasiodiplodin. Similarly, the node with *m/z* 275.165 represents the [M- H_2_O+ H]^+^ adduct of **18**. The node with *m/z* 293.139 (**19**) is adjacent to **17** and has a high similarity to the [M + H]^+^ adduct of dihydroresorcylide. The node with *m/z* 291.16 (**20**) could match the [M + H]^+^ adduct of (E)-9-etheno-de-O-methyllasiodiplodin or 12E,15R-5-hydroxy-3-methoxy-16-methyl-8,9,10,11,14,15-hexahydro-1H-benzo[c][1]oxacyclodocecin-1-one, with a different double bond position in the main ring. Based on the observed MS2 fragments, including fragments with *m/z* 205.0868, *m/z* 191.0707, *m/z* 177.0917, and *m/z* 149.0602, this node possibly represents (E)-9-etheno-de-O-methyllasiodiplodin. SNAP-MS also indicated the existence of one more hydroxyl group in the main ring, which contributed to the identification of an [M + H]^+^ adduct of compound **21** consisting of planar structures of 7,12-dihydroxy-14-methoxy-3-methyl-3,4,5,6,7,8,9,10-octahydro-1H-benzo[c][1]oxacyclododecin-1-one, 5,12-dihydroxy-14-methoxy-3-methyl-3,4,5,6,7,8,9,10-octahydro-1H-benzo[c][1]oxacyclododecin-1-one, or 4,12-dihydroxy-14-methoxy-3-methyl-3,4,5,6,7,8,9,10-octahydro-1H-benzo[c][1]oxacyclododecin-1-one. Due to the presence of some fragment peaks, including peaks with *m/z* 233.1181, *m/z* 195.1017, *m/z* 193.0863, and *m/z* 69.0701, compound **21** was identified as 7,12-dihydroxy-14-methoxy-3-methyl-3,4,5,6,7,8,9,10-octahydro-1H-benzo[c][1]oxacyclododecin-1-one. In addition, nodes with *m/z* 249.15, *m/z* 277.1810, and *m/z* 357.1380 are separately predicted to be [M- H_2_O+ H]^+^ adducts of [4]-gingerol (**22**), [6]-gingerol (**23**), and 4-(5-hydroxy-3-oxodecyl)-2-methoxyphenyl hydrogen sulfate (**24**), with -C_2_H_4_ and -SO_3_ edge differences between them. The nodes with *m/z* 305.1140 and *m/z* 277.1190 are predicted to be 2-(3-carboxypropanoylamino)-3-indol-3-ylpropanoic acid (**25**), and PyroGlu-Phe (**26**), respectively.

Note S2. Other biosynthetic potentials of *Cosmosporella* sp. VM-42.

The BGC in region 14.2 encodes a non-reducing T1PKS, and KnownClusterBlast reveals that 44% of genes in a hex-pk23 polyketide cluster from *Endocarpon pusillum Z07020* and 21% of genes in a duclauxin cluster from *Talaromyces stipitatus* ATCC 10500 match to the query sequence. The predicted core PKS shares 57% amino acid sequence identity with that in *E. pusillum Z07020*, possessing a biosynthetic unit of SAT-KS-AT-PT-ACP-ACP-TE (Figure S11A). By heterologous protein expression in the HEx synthetic biology platform, the core enzyme in *E. pusillum Z07020* was demonstrated to produce two naphthalene pyrones, including prephenalenone and its dehydrated compound 6,7,9-trihydroxy-3-methyl-1H-benzo[f]chromen-1-one (Harvey et al., 2018). Unlike the duclauxin gene cluster, our BGC does neither encode a flavin-dependent monooxygenase that catalyses the C2 aromatic hydroxylation of prephenalenone and opening of the γ-pyrone ring simultaneously, nor a P450 monooxygenase that participates in the subsequent redox modifications and dimerization for the formation of duclauxin (Gao et al., 2018, 2016). A predicted cupin protein encoded in our BGC shares 63% identity with a cupin family dioxygenase DUXM in *T. stipitatus* ATCC 10500 that performs an oxidative cleavage of the peri-fused tricyclic phenalenone and affords a transient hemiketal-oxaphenalenone intermediate (Gao et al., 2018). However, due to the lack of the phenalenone substrate reported in the literature, this enzyme may be responsible for a different reaction in *Cosmosporella* sp. VM-42 and lead to different products, which remains to be further determined.

Region 11.3 shares similarity with an azasperpyranone A BGC in *Aspergillus terreus* NIH2624, encoding two highly reducing PKS genes (48% amino acid sequence identity, KS-AT-DH-cMT-ER-KR-ACP; 52% amino acid sequence identity, KS-AT-ACP-cMT-TD), as displayed in Figure S11B. Furthermore, the putative fungal Zn(2)-Cys(6) transcription factor and the EmrB/QacA drug resistance transporter also showed 31% and 47% amino acid sequence identity with their counterparts *A. terreus* NIH2624. It is important to note that azapyranone A is not biosynthesized by enzymes encoded in this *A. terreus* NIH2624 BGC alone, but that it is synthesized synergistically by four core enzymes, two non-reducing PKS, one highly reducing PKS, and one NRPS-like, encoded in two separate clusters. The expression of the respective genes is coordinated by collaborative transcriptional regulation based on three regulators in the two clusters (Huang et al., 2020). Whether there are similar synergistic biosynthetic pathways in *Cosmosporella* sp. VM-42 is worthy of further exploration.

Region 9.1 includes a T1PKS BGC sharing partial similarities with a betaenone A BGC from *Phoma betae* (BGC0001264) and a calbistrin A biosynthetic gene cluster from *Emericella variecolor* (BGC0002168), respectively. The core genes of these two reference BGCs encode highly reducing PKS units with the domain organization of KS-AT-DH-cMT-inactive enoylreductase (ER^0^)-KR-ACP-R, which act together with a trans-acting ER to form a decalin scaffold, as shown in Figure S11C (Tao et al., 2021; Ugai et al., 2015). Interestingly, this polyketide synthase can also participate in the production of the polyene portion of calbistrins with the co-operation of the trans-acting C-methyltransferase CALH (Figure S11C) (Tao et al., 2021). Compared to these known BGCs, the core enzyme encoded in region 9.1 is unexpectedly divided into three genes, encoding a cMT-KR-ACP polypeptide, a standalone DH domain, and a standalone KS domain, without an AT domain. This also raises other questions about *Cosmosporella* sp. VM-42. For example, whether this core protein can form the decalin structure in the absence of an acyltransferase domain or whether it needs the help of other enzymes encoded in this cluster. Furthermore, the tailoring enzymes encoded in this BGC are different from the known BGCs and may contribute to the formation of different compound types than betaenone A and calbistrin A. Gene ontology analysis also indicates that one gene in this region encodes an Snf2-related domain with ATP-dependent chromatin remodelling activity. Snf2 proteins represent a family of helicase-like proteins that direct energy derived from ATP hydrolysis into the mechanical remodelling of chromatin structure (Ryan and Owen-Hughes, 2011). However, there is currently no literature on the role of SNF2 domains in filamentous fungi and their roles in fungal secondary metabolites production.

Region 7.2 is a hybrid BGC encoding NRPS and T1PKS. Although KnownClusterBlast returned high similarity with a UNII-YC2Q1O94PT BGC from *Alternaria alternata* (BGC0001252), we noticed that the reference BGC only has the annotation for the PKS-encoding gene and is missing other genes. Furthermore, the NPRS, encoded by the core gene in region 7.2, is predicted to use Ival as substrate, which is not a (known) intermediate in UNII-YC2Q1O94PT biosynthesis. Therefore, the final product of the NRPS and T1PKS encoded by region 7.2 remains still unknown.

Region 7.1 includes an NRP BGC with genes matching 62% of genes in the metachelin C BGC from *Metarhizium robertsii ARSEF 23* (BGC0002710), which can produce siderophores. In Region 13.1 two genes encoding an NRPS and a cytochrome P450 have significant matches with genes in the gliovirin BGC from *Trichoderma virens Gv*29-8 (BGC0001609).
